# Extra-cellular matrix remodeling as a unique mechanism of expansion of periprostatic adipose tissue: a potential driver of prostate cancer aggressiveness

**DOI:** 10.1101/2023.01.05.522843

**Authors:** David Estève, Aurélie Toulet, Mathieu Roumiguié, Dawei Bu, Mathilde Lacombe, Sarah Pericart, Chloé Belles, Cécile Manceau, Cynthia Houël, Manuelle Ducoux-Petit, Nathalie Van Acker, Stéphanie Dauvillier, Yiyue Jia, Marine Hernandez, Mohamed Moutahir, Camille Franchet, Nicolas Doumerc, Mathieu Thoulouzan, Sophie Le Gonidec, Philippe Valet, Bernard Malavaud, Odile Burlet-Schiltz, Anne Bouloumié, Philipp E. Scherer, Delphine Milhas, Catherine Muller

**Affiliations:** Institut de Pharmacologie et de Biologie Structurale (IPBS), Université de Toulouse, CNRS, Université Toulouse III – Paul Sabatier (UPS), Toulouse, France. Équipe Labélisée Ligue Nationale contre le Cancer.; Department of Urology and Renal Transplantation, Toulouse University Hospital; Institut Universitaire du Cancer de Toulouse, Oncopole, Toulouse, France.; Touchstone Diabetes Center, Department of Internal Medicine and Department of Cell Biology, University of Texas Southwestern Medical Center, Dallas, TX, USA.; Department of Pathology, Institut Universitaire du Cancer de Toulouse, Oncopole, Toulouse, France.; Institut des Maladies Métaboliques et Cardiovasculaires, Université de Toulouse, INSERM, UPS, Toulouse, France; Infrastructure Nationale de Protéomique, ProFI, FR 2048, Toulouse, France; Institut RESTORE, Université de Toulouse, CNRS U-5070, EFS, ENVT, INSERM U1301, UPS, Toulouse, France.

## Abstract

One of the most striking features of the adipose depot surrounding the prostate (periprostatic adipose tissue, PPAT) is that its accumulation is independent of body mass index. Its volume varies considerably between individual with some patients exhibiting abundant PPATs that have been correlated to occurrence of aggressive prostate cancer (PCa). However, abundant PPAT are not defined at biological levels. We used a new statistical approach to define abundant PPAT by normalizing PPAT volume to prostate volume in a cohort of 351 patients with a linear regression model. Applying this definition, we confirmed the link between abundant PPAT and PCa aggressiveness, therefore validating our approach. At biological levels, we showed that abundant PPAT exhibited extensive extracellular matrix remodeling, notably of the collagen network, decreasing the mechanical constraints in hypertrophic adipocytes leading to an inflammation free-expansion. Degradation of the most abundant collagen in AT, collagen VI was associated with increased production of endotrophin, a signaling peptide derived from AT, that was also elevated in the urine of patients with abundant PPAT confirming the clinical relevance of our results. These results highlight a unique mechanism of expansion of an adipose depot and open new mechanistic avenues to explain its role in prostate-related disorders.

## Introduction

In addition to the major subcutaneous (SC) or visceral white adipose tissues (WAT), smaller and organ-associated adipose depots are dispersed throughout the body such as mammary, epicardial, perivascular and bone marrow adipose tissues (AT) (1). These AT could be defined as “organ-associated” AT and their primary function could be to regulate proximal organ function rather than participating to general metabolism homeostasis (1). Accordingly, it is expected that these AT will exhibit changes in their tissular organization, responses to external stimuli as well as metabolic and secretory functions in comparison to “classical” adipose depots. Periprostatic AT (PPAT) is a WAT which surrounds the prostate. Although it shares common vasculature with the prostate, it is separated from it by a fibro-muscular sheet called the prostate capsule. Nevertheless, one-third of the anterior zone of the prostate is in direct contact with this tissue (2). Like other WAT, PPAT comprises mainly adipocytes and other cell types in the stromal vascular fraction (SVF), including adipose progenitor cells, fibroblasts, endothelial and immune cells, all of which are embedded in the extracellular matrix (ECM) (3,4). Also, as a WAT, PPAT is an active metabolic and secretory organ (1). Accordingly, PPAT can impact various prostate related diseases including benign prostate hyperplasia (BPH), erectile and urethral dysfunctions and prostate cancer (PCa) although its physiological roles remain mainly uncharacterized (5–7).

One of the most striking features of PPAT is that its accumulation is independent of body mass index (BMI) that is, to our knowledge, a unique behaviour among adipose depots (6,8). By comparison with abdominal pelvic adipose tissue (APAT), which behaves like a typical WAT, we found that PPAT (in both healthy and cancer-bearing patients) has a sparse vascular network responsible for a chronic hypoxic state leading to a dense collagen network typical of fibrosis in lean patients (8). In opposition to the control AT, no increase in this collagen network was observed in PPAT between lean individuals and individuals with obesity (8). We proposed that this obesity-like organization of the ECM in PPAT does not adapt to external signals but provides an important physical constraint on PPAT expansion (8). A recent study demonstrated that obesity was not associated with adipocyte hypertrophy in PPAT (9) consistent with the idea that PPAT is in a state of maximum expandability in most individuals.

However, the volume of PPAT considerably varies between patients, highlighting that PPAT has the capacity to expand upon uncharacterized mechanisms (8,10,11). Recent clinical studies have demonstrated that patients with large amounts of PPAT have more aggressive PCa (6). PPAT abundance correlates with prostate tumors with high Gleason scores and local and distant dissemination (6,10–16). The role of abundant PPAT on other prostate-related disorders has not been yet investigated. The biological mechanisms that result in expansion of PPAT are still unknown. In this study, we first developed a more refined definition of abundant PPAT that takes into account the correlation between PPAT and prostate volume demonstrated in previous studies (17,18). Using this definition, we validated that PPAT abundance was correlated to PCa aggressiveness as described in previous studies (6,10–16) and used these validated samples to characterize their biological characteristics. We showed that abundant PPAT exhibited extensive ECM remodeling, notably of the collagen network, decreasing the mechanical constraints in hypertrophic adipocytes leading to an inflammation free-expansion. Degradation of the most abundant collage in AT, collagen VI (COL6) was associated with increased production of endotrophin (ETP), a signaling peptide derived from AT (19). We further confirmed that ETP was elevated in the urine of patients with abundant PPAT showing that ECM remodeling exists at clinical level. This first functional and structural biological characterization of PPAT points to an unusual mechanism of AT expansion based on the reshaping of the ECM and opens new avenues to explain its role in prostate disorders.

## Results

### A new statistical approach to define abundant PPAT

As PPAT volume is reported to correlate with prostate volume, recent studies have used the ratio of PPAT volume to prostate volume in an individual – the normalized periprostatic fat volume (NPFV) – to compare PPAT abundance between individuals in order to better define the relative fat volume in the periprostatic area (17,18). To improve this definition, we developed a measure that expresses the volume of PPAT with respect to the volume of the prostate normalized to a population rather than to an individual. We first used a cohort of 351 patients awaiting prostatectomy for PCa by using pre-operative multiparametric magnetic resonance imaging (mpMRI), from which we measured prostate and PPAT volume within well-defined anatomical limits (Table S1 and Fig. 1A). We found a strong correlation between the volume of the prostate determined by mpMRI before prostatectomy and its weight after prostatectomy (Fig. 1B), validating the accuracy of the measurement method. A group of 20 patients that underwent mpMRI for non-oncological pathologies were also included in the study. In all patients (with or without cancer), PPAT volume was independent of BMI (Fig. 1C) and correlated with prostate volume (Fig. 1D). As expected, both SC-AT area (Fig. S1A) and perirectal AT area (Fig. S1B) correlated with BMI in the cohort. Based on this strong correlation between PPAT volume and prostate volume, we modeled the expected PPAT volume according to the prostate volume by using a linear regression model and calculated PPAT abundance with the residual values from this model. The residual values represent the difference between the volume of PPAT quantified by mpMRI and the amount of PPAT predicted by the model (Fig. 1E). The cohort was divided into quartiles according to residuals values allowing to identify patients with the least PPAT in the 1^st^ quartile (low PPAT) and patients with the most PPAT in the 4^th^ quartile (high PPAT) (Fig. 1 E-F). Patients in the 2^nd^ and 3^rd^ quartile were classified as having a mid PPAT abundance (Fig. 1 E-F). No differences were observed between patients in the four quartiles when we analyzed prostate volume (Fig. 1G), BMI (Fig. 1H) or age (Fig. 1I).

**Figure 1:**
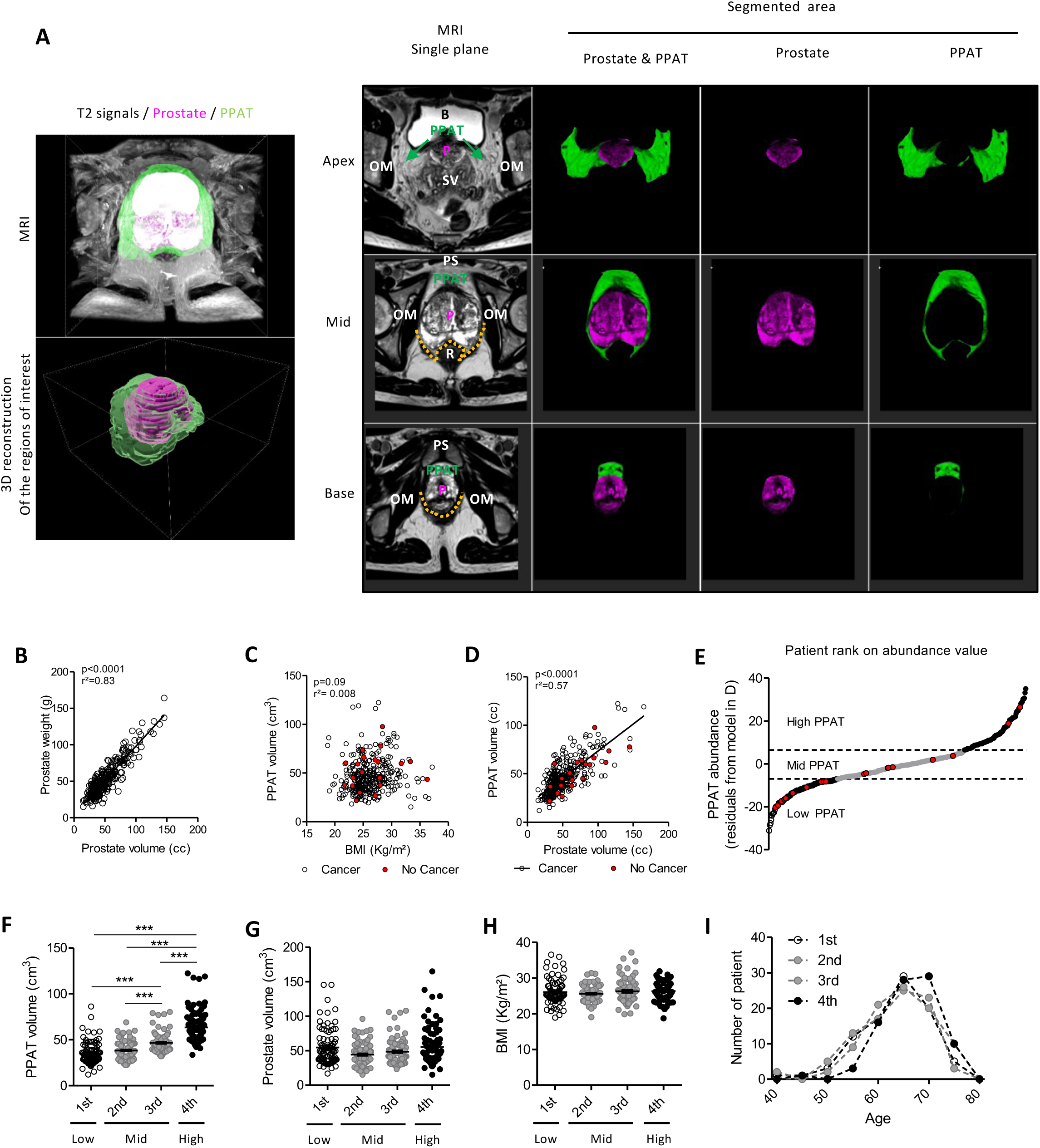
A new statistical approach to define abundant PPAT relative to prostate volume. **A)** Left panel: Representative 3D projection and reconstruction of axial sections from T2-weighted mpMRI signals and regions of interest delineating the prostate (pink) and the PPAT (green); Right panel: single axial MRI sections from the base, middle, and apex of the prostate. The localization of PPAT, prostate (P) and the anatomical limits of PPAT are shown on single axial MRI section (B, Bladder; OM, Obturator Muscle; P, Prostate; PPAT, Periprostatic Adipose Tissue; PS, Pubic Symphysis; R, Rectum; SV, Seminal Vesicles; orange dotted lines indicate Denonvillier’s fascia). Segmented area represents the region of interest delineating the prostate (pink) and the PPAT (green) for each single MRI section presented. **B)** Correlation between prostate weight recorded after prostatectomy and prostate volume measured on mpMRI (n=351). Linear regression was used to draw the slope of best fit and linear correlation coefficient (r^2^) and P-value are indicated (p). **C)** Lack of correlation between prostate volume and the BMI of the patient (n=371); patients without PCa are indicated in red. **D)** Correlation between PPAT and prostate volume (n=371); patients without PCa are indicated in red. Linear regression was used to draw the slope of best fit and linear correlation coefficient (r^2^) and P-value are indicated (p). **E)** Scatter plot of patients ranked according to PPAT abundance showing patients with low, mid and high PPAT abundance. PPAT abundance was calculated from the residual value for each patient with respect to the linear regression model in part D (n=371); patients without PCa are indicated in red. **F)** PPAT volume measured on MRI according to the abundance quartile (n=86-87 per quartile). **G)** Prostate volume of the patients in each abundance quartile (n=86-87 per quartile). **H)** BMI of the patients in each abundance quartile (n=86-87 per quartile). **I)** Distribution of patients’ age in each abundance quartile (n=86-87 per quartile). Individual patient’s data are indicated by dots, bars indicate means ± SEM. Statistical differences between groups were evaluated by ANOVA followed by a post-test for linear trend (*** p< 0.001).

We then validated in our cohort (whose clinical and biological parameters are presented in Table S2) the relationship between PPAT abundance, as defined above, and PCa aggressiveness. Of note, this cohort included only patients who underwent prostatectomy, since the structural and functional characterization of abundant compared to low PPAT was the goal of our study. Therefore, the cohort was rather homogenous in the aggressiveness of their cancers, according to prostatectomy indications (20). Most patients in our PCa cohort were graded 2 or 3 on the ISUP scale (Table S2). Nevertheless, the proportion of patients with an ISUP grade ≥ 3 increased with PPAT abundance and was significantly different between the low PPAT and high PPAT groups (Fig. 2A and Table S3). In addition to the ISUP score, the percentage of high-grade lesions in the tumors of patients in the high PPAT group was significantly greater than in the patients in the low PPAT group (Fig. 2B and Table S3). Moreover, there was a significant association between the abundance of PPAT and the concentration of prostate-specific antigen in the blood of patients with PCa prior to surgery (Table S3). Multivariate analysis using a logistic regression model found that PPAT abundance was an independent risk factor for PCa with an ISUP score ≥ 3 (Fig. 2C). Therefore, these results in accordance with the literature (6,10–16) validated our ranking of PPATs abundance. We then characterized its biological characteristics.

**Figure 2:**
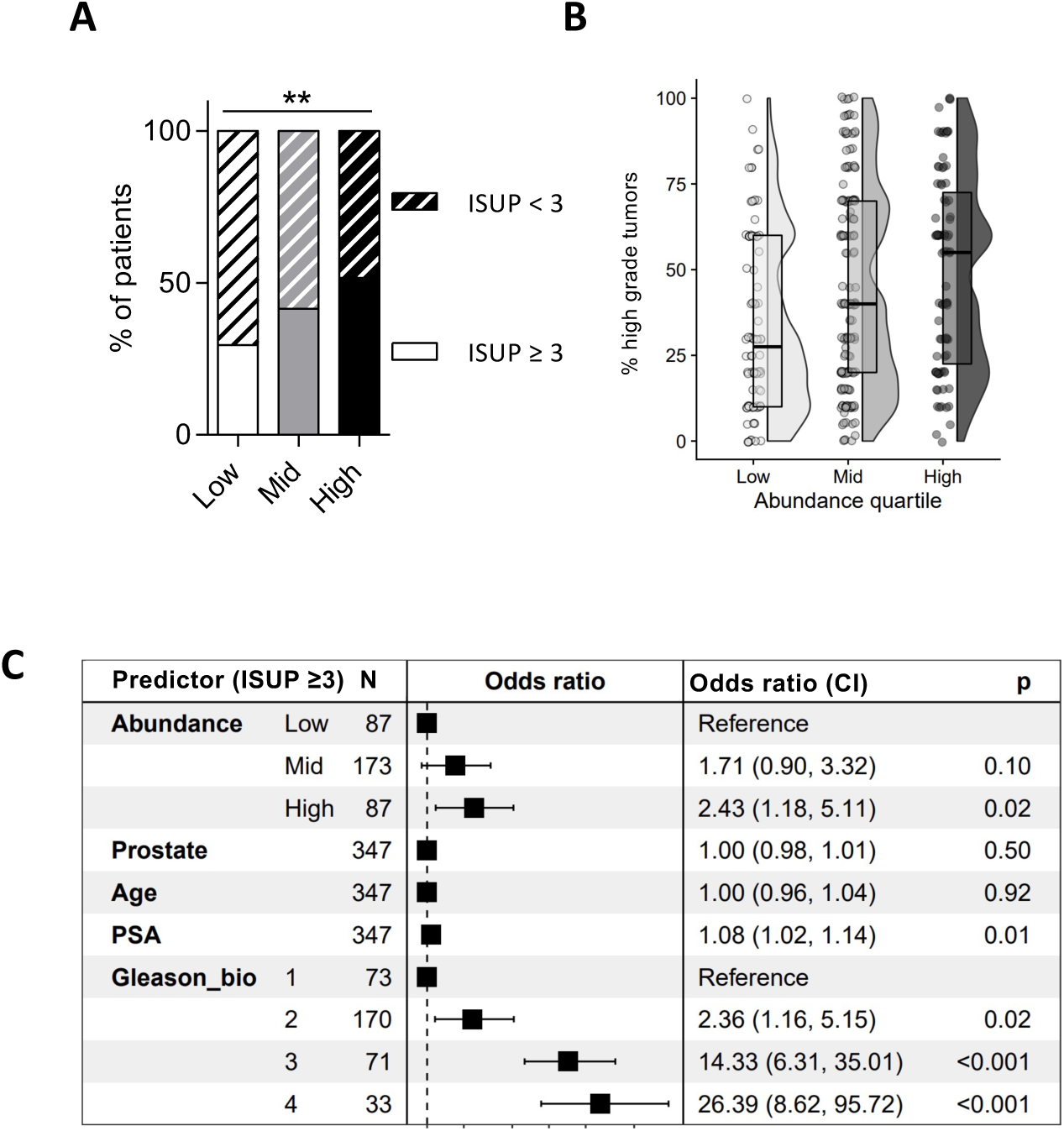
Abundant PPAT is associated with aggressive PCa. **A)** Percentage of patients in the low, mid and high PPAT groups who have ISUP scores < 3 and ≥ 3. **B)** Raincloud plot of the percentage of high grade tumors observed in prostate tumor slices from patients in the low, mid and high PPAT groups (Low, n= 88; Mid, n=176; High, n=87). **C)** Multivariate analysis using logistic regression modelling of the effect of the indicated variables on the relative risk (odds ratio) that the PCa has an ISUP score ≥ 3 (Abundance, low, mid and high PPAT groups; Prostate, prostate volume (cm^3^); Age (years); PSA, plasma PSA concentration (ng/ml); Gleason_bio, Gleason score on biopsy). Significant differences between groups were evaluated by Chi-square test for qualitative variables (A), by using ANOVA with post-test for linear trend (**p< 0.01; B). Odds ratio were determined by using the multivariate logistic regression model (C).

### PPAT expands by adipocyte hypertrophy with no associated inflammation

AT expands in response to surplus energy by two main mechanisms: adipocyte enlargement, or hypertrophy, and adipocyte hyperplasia, in which new adipocytes are recruited by adipogenesis from progenitors (21). The balance between hypertrophy and hyperplasia depends on the adipose depot concerned (21). Paraffin-embedded PPAT sections stained with HE were used to measure the diameters of the adipocytes by a semi-automated method, Adiposoft^©^(Fig. 3A). The adipocyte size distribution curve was shifted towards larger cells in PPAT from patients in the high PPAT group when compared with those in the low PPAT group, and the mean adipocyte diameter was larger (Fig. 3B), indicating adipocyte hypertrophy. To quantify adipose progenitors in the SVF, cells were immunostained with antibodies against the markers CD45, CD34 and CD31 and analyzed by flow cytometry. The percentage of cells in this fraction that were progenitor cells (defined as CD45-/CD34+/CD31-cells) was similar in PPAT from patients in the various groups (Fig. 3C). Furthermore, when additional markers (MSCA1 and CD271) were used to characterize the sub-populations of adipose progenitors (22), the proportions of the various progenitor subsets (the adipogenic MSCA1+; myofibrogenic MSCA1-/CD271+, and immature MSCA1/CD271-subsets), were similar in all groups (Fig. S2A), indicating no increase in progenitors in abundant PPAT. We conclude that adipocyte hypertrophy rather than hyperplasia is the mechanism of PPAT expansion in patients with abundant PPAT.

**Figure 3:**
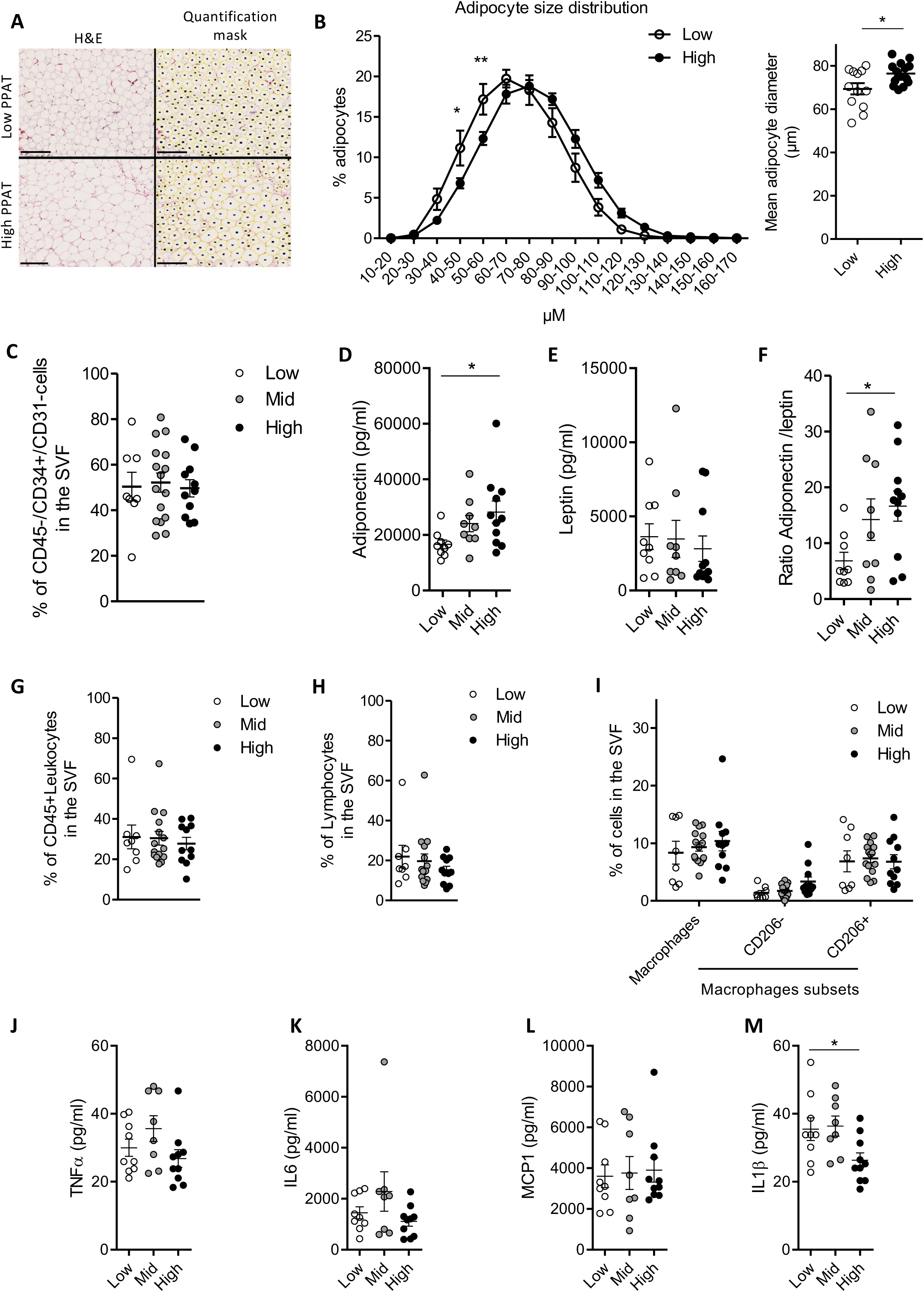
PPAT expands by adipocyte hypertrophy with no associated inflammation. **A)** Representative images of PPAT sections stained with hematoxylin and eosin (H&E) and used to quantify adipocyte diameter in PPAT from patients in the low and high groups. The corresponding quantification mask is shown. Scale bars, 250μm. **B)** The size distribution (left panel) and mean diameters (right panel) of adipocytes in PPAT from patients in the low (n=12) and high (n=14) groups. **C)** Percentage of progenitor cells (CD45-/CD34+/CD31-) in the stroma-vascular fraction (SVF) of PPAT from patients in the low (n=8), mid (n=16) and high (n=11) groups, as quantified by flow cytometry quartile). **D-F)** Quantification of the concentration of adiponectin (D) and leptin (E) in the conditioned medium of PPAT from patients in the low (n=9), mid (n=9) and high (n=11) groups, and the adiponectin to leptin ratio (F). **G)** Percentage of cells in the SVF of PPAT from patients in the low (n=8), mid (n=15) and high (n=11) groups that are CD45+ leukocytes. **H**) Percentage of cells as in (G) that are low FSC/low SSC/CD45+/CD3+ T cells in the low (n=8), mid (n=15) and high (n=11) groups. **(I)** Percentage of cells as in (G) that are CD45+/CD14+ macrophages (all macrophages), CD45+/CD14+/CD206-recruited macrophages (CD206-), or CD45+/CD14+/CD206+ resident macrophages (CD206+) in the low (n=8), mid (n=15) and high (n=11) groups. All percentages in G-I were quantified by flow cytometry. **J-M)** Quantification of the concentrations of TNFα **(J)**, IL6 **(K)**, MCP1 **(L)** and IL1β **(M)** in the conditioned medium of PPAT from patients in the low, mid and high groups (n=8-10). Bars indicate means +/-SEM. Two-way ANOVA followed by Bonferroni post-test was used to analyze adipocyte diameter. Other statistical differences were evaluated by ANOVA with post-test for linear trend (*p<0.05; ** p<0.01).

Adipocyte hypertrophy during AT expansion is usually associated with changes in secretion related to AT dysfunction (3,4). Key among these changes is increased secretion of leptin and, in parallel, decreased secretion of adiponectin (3,4). The decrease in the ratio of adiponectin/leptin secretion has been proposed as a marker of dysfunctional AT in metabolic disorders and inflammation (3,4). Despite the adipocyte hypertrophy we saw in PPAT from patients in the high PPAT group, more adiponectin was secreted by this PPAT than by PPAT from patients in the low PPAT group (Fig. 3D), whereas leptin secretion by both tissues was similar (Fig. 3E). Accordingly, the adiponectin/leptin secretion ratio was greater in high than in low PPAT (Fig. 3F). Thus, the adipocyte hypertrophy observed in high PPAT is not associated with the changes in secretion that are usually observed in dysfunctional AT.

The proportion of immune system cells in AT generally increases as the tissue expands and this is associated with the onset of chronic low-grade inflammation (23). We measured the proportion of immune cells in PPAT from patients in the three groups by immunostaining the cells in the SVF with antibodies against CD45 and other specific markers of immune population subsets and analyzed them by flow cytometry. The proportion of CD45+ leukocytes in this fraction was similar in the various groups of patients (Fig. 3G), as was the proportion of total lymphocytes (defined as CD45+/low side scatter) (Fig. 3H) and of various lymphocyte subsets (Figure S2B). The proportion of macrophages (defined as CD45+/CD14+ cells), including the sub-populations that express or do not express CD206, were also similar in the various groups (Fig. 3I). These data indicate that PPAT expansion is not associated with increased immune cell infiltration.

To characterize further the inflammatory state of abundant PPAT, we quantified secretion of several proinflammatory cytokines by PPAT from patients in the various groups. No significant differences were seen between the amounts of TNF-α, IL6, or MCP-1 secreted (Fig. 3 J-L), and the tissue from patients in the high PPAT group secreted slightly less IL1β than the others (Fig. 3M). Thus, the PPAT from patients in the high PPAT group does not secrete more proinflammatory cytokines than the PPAT from the patients in the mid and low PPAT groups. Together, these data indicate that although PPAT expands by adipocyte hypertrophy, this expansion does not cause the inflammation observed in other ATs that expand by this mechanism.

### Under-representation of mechano-sensing proteins in adipocytes from abundant PPAT

According to our findings described above, adipocytes are the main cell type that changes in the PPAT from patients in the high PPAT group. Adipocytes from high PPAT and low PPAT groups were isolated and analyzed by proteomic approaches. Principal component analysis of the proteomic dataset separated the patients according to the abundance of their PPAT (Fig. 4A). Component 1, which separated the patients according to their PPAT groups, explained 52% of the variance in the proteomic dataset (Fig. 4A). Whereas the low PPAT patients grouped closely on the scatter plot, those with high PPAT spread widely (Fig. 4A). Of the 4033 proteins identified by proteomic, statistical analysis found 352 whose abundance differed in PPAT from patients in the high and low PPAT groups: 157 were over-represented and 195 were under-represented in the samples from the high PPAT group when compared to those from the low PPAT group (Fig. 4B). Hierarchical clustering by using Ward’s method on differentially represented proteins grouped the samples according to their abundance status (Fig. 4C).

**Figure 4:**
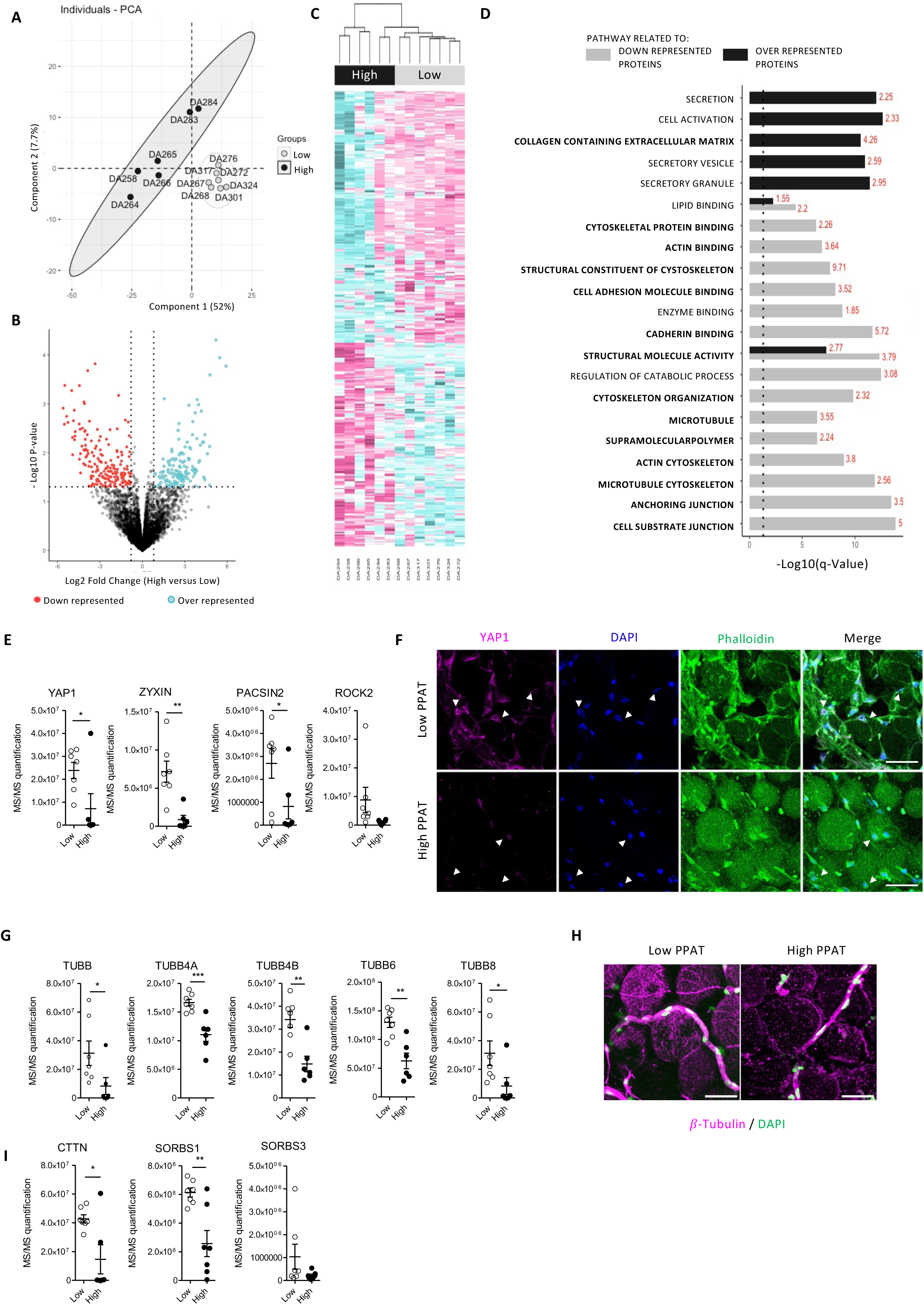
Under-representation of mechano-sensing proteins in adipocytes from abundant PPAT. **A)** Principal component analysis of the proteomics datasets from adipocytes isolated from the PPAT of individual patients (each indicated by a dot and a label) in the low and high groups (n= 6-7 per group). The ellipses indicate grouping according to PPAT abundance. **B)** Volcano plot representing the log2 fold-change of the relative protein representation in the proteomics dataset and the –log10 of the p-value calculated by using limma differential expression analysis (n=6-7 per group), showing under-represented (pink) and over-represented (green) proteins. **C**) Hierarchical clustering of differentially under-represented (pink) and over-represented (green) proteins in the PPAT of patients in the low (grey) and high (black) quartiles. **D)** Cellular components and biological functions of the over-represented (black) and under-represented (grey) proteins in the PPAT of patients in the high quartile, as determined by gene ontology enrichment terms. The –log10 of the q-Value are represented in x axis and the percentage of proteins that were differentially expressed in the pathways are mentioned in red on the side of each bar plot. **E)** Quantification of the indicated proteins in the PPAT of patients in the low and high quartile (n=6-7 per group) by mass spectrometry. **F)** Whole PPAT from patients in the low and high groups were stained with anti-YAP1 antibody and phalloidin (to stain actin and facilitate cell visualization) and DAPI (to stain DNA). White arrows indicate nuclei of adipocytes. Representative maximum intensity projection of images obtained by 3D confocal microscopy are shown. Scale bars, 50 μm. G) Quantification of the indicated isoforms of tubulin, as in E (n=6-7 per group). **H)** Whole PPAT from patients in the low and high groups were stained with anti-ß-tubulin antibody and DAPI, as in F. **I)** Quantification of the indicated proteins, as in E (n=6-7 per group). Bars indicate means ± SEM. Statistical differences were calculated by using Student’s T test. *p < 0.05; ** p< 0.01; *** p<0.001.

To further characterize the differences between adipocytes in the PPAT of patients in the high and low PPAT groups, we used gene ontology enrichment analysis of the proteomic dataset to identify cellular components and molecular functions. We found large differences in the abundance of proteins involved in cytoskeletal organization, cell-ECM interactions and, to a lesser extent, lipid metabolism (Fig. 4D). From this analysis, the main characteristics of adipocytes from abundant PPAT can be described as having, foremost, an under-representation of proteins involved in mechano-transduction and sensing stiffness (ontology terms in bold in Fig. 4D). Among them, YAP1 (Yes-associated protein 1) is the most under-represented protein in the PPAT of patients in the high PPAT group (log2 fold-change = –5.8) when compared with the low PPAT group (Fig. 4E). YAP1 is a key mechano-transducer that senses mechanical stimuli and relays them to regulate transcription and, thus, many aspects of cell behavior (24). ZYXIN (25) and PACSIN2 (26), proteins that participate in the nuclear translocation and activation of YAP1 upon mechanical stress, were also significantly under-represented in the most abundant PPAT, as was ROCK2, a target of the YAP/TEAD transcriptional complex (24) (Fig. 4E). We confirmed by immunofluorescence microscopy that PPAT from patients in the high PPAT group contained much less YAP1 than that from patients in the low PPAT group (Fig. 4F).

The second important characteristic of adipocytes from abundant PPAT identified by pathway enrichment analysis was under-representation of several tubulin isoforms (Fig. 4G), which was accompanied by altered microtubule organization seen by immunofluorescence microscopy (Fig. 4H). Proteins involved in cytoskeletal organization, such as cortactin (CTTN), an actin nucleation factor that influences the stability of actin filaments, and members of the vinexin family (SORBS-1 and SORBS-3), which are involved in contractile force generation in response to mechanical stress (27) were also under-represented (Fig. 4I).

A third notable characteristic of adipocytes from the high PPAT group was the abundance of proteins involved in lipid metabolism (Fig. 4D), which may explain, at least in part, the observed adipocyte hypertrophy. Several key proteins involved in lipolysis were under-represented in PPAT from patients in the high PPAT group, including adipocyte triglyceride lipase (encoded by *PNPLA2*), hormone-sensitive lipase (encoded by *LIPE*), perilipin1 (encoded by PLIN1) and Abhydrolase domain-containing protein 5 (encoded by ABHD5) (Fig. S3A). For PNPLA2 and LIPE, a downward trend regulation at mRNA levels according to abundance status was confirmed by RT-qPCR, whereas the level of MGLL (encoding Monoacylglycerol Lipase) was unchanged (Fig. S3B). By contrast, adiponectin (encoded by ADIPOQ) and several lipid transporters that take up fatty acids like CD36 or FABP4 (Fatty Acid Binding protein 4) were over-represented (Fig. S3A). Regulation of their mRNA levels according to abundance status was confirmed by RT-qPCR and similar differences were also observed for the Long-chain fatty acid transport protein 4 (encoded by SLC27A4) (Fig. S3C).

### The collagen network in abundant PPAT is relatively loose

The under-representation of proteins involved in mechano-sensing and cytoskeletal contractile force generation in abundant PPAT suggest that these PPAT might be subject to fewer mechanical constraints by ECM than less abundant PPAT, which may account for its expansion without inflammation. To investigate this hypothesis, we stained the collagen fibers with picrosirius red, imaged the staining by 3D confocal microscopy and reconstructed images of the fiber network in 3D. In the reconstructions, the signal from collagen fibers was weaker in PPAT from patients in the high PPAT group than it was in the patients in the low PPAT group (as shown by the maximum intensity projection), and the collagen network was substantially less dense and had thinner fibers (Fig. 5A). These observations were verified by using Imaris software to quantify the volume and length of the collagen fibers as well as the total number of branches seen in the 3D reconstructions (Fig. 5 B–D). To further confirm our findings, we used a different approach based on quantification of hydroxyproline, a component of collagen that comprises around 13.5% of its amino acid composition. Consistent with the imaging analysis of the collagen fiber network, we found significantly less hydroxyproline in PPAT from patients in the high PPAT group than in patients in the low PPAT group (Fig. 5E). Together, these analyses indicate that abundant PPAT has a looser collagen network than less abundant PPAT, supporting the hypothesis that mature adipocytes in abundant PPAT are subject to fewer mechanical constraints than those in less abundant PPAT.

**Figure 5:**
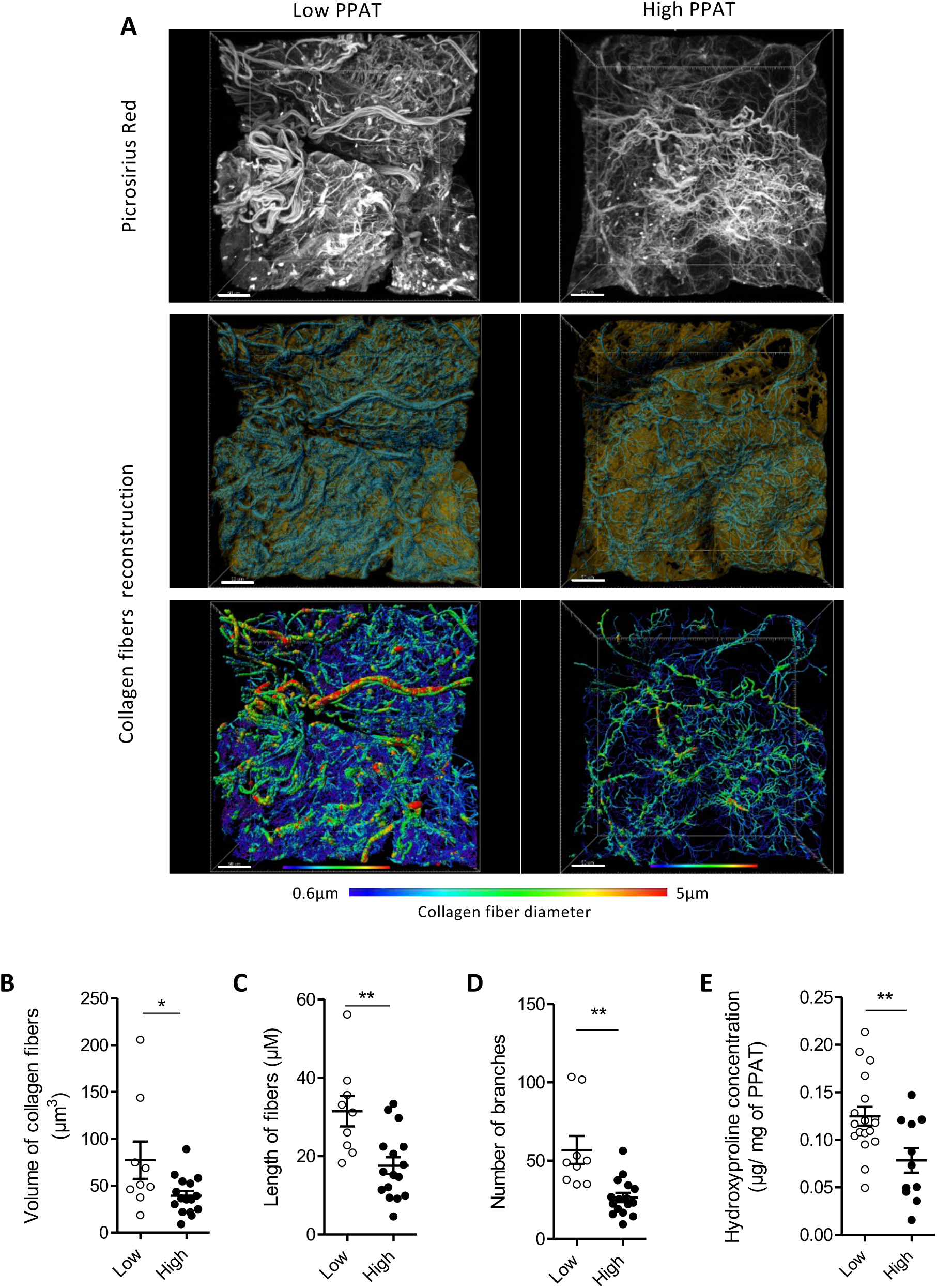
The collagen network in abundant PPAT is relatively loose. **A)** Whole PPAT from representative patients in the low and high groups were stained with picrosirius red to show collagen fibers, imaged in 3D by confocal microscopy (top), reconstructed in 3D (middle), and the reconstructed collagen fiber images were isolated and color-coded according to their diameter (bottom). Scale bars, 50 μm. **B-D)** Various morphometric parameters were quantified in 3D images of PPAT from patients in the low and high groups after collagen fiber reconstruction (n=9-16): total collagen volume (B), length of collagen fibers (C), and number of branches per collagen fiber (D). **E)** Quantification of hydroxyproline concentration in PPAT from patients in the low (n=18) and high groups (n=11). Bars indicate means ± SEM. Statistical differences were calculated by using Student’s T test. *p < 0.05; ** p< 0.01.

### Abundant PPAT exhibits increased collagen degradation likely by MMP9

Our findings indicate that abundant PPAT contains less collagen organized in a looser network than in less abundant PPAT. To determine whether this reflects less collagen synthesis or more degradation of the proteins, we used RT-qPCR to compare the expression of the genes encoding the major fibrillar collagens (*COL1A1* and *COL3A1*) as well as the 3 different alpha chains of the most abundant non-fibrillar collagen, collagen VI (28) (*COL6A1*, *COL6A2* and *COL6A3*) in PPAT from patients in the high and low PPAT groups. We saw no differences between them (Fig. 6A) suggesting that the smaller amount of collagen and looser network in abundant PPAT is not due to less synthesis but may be due to increased degradation. To investigate this hypothesis, we stained PPAT samples with a fluorescent peptide (F-CHP) that binds specifically to denatured collagen strands (29) and imaged the staining by 3D confocal microscopy. The images showed a punctate staining located alongside fibrillar structures in PPAT from patients in the high PPAT group, whereas very weak signals were seen in PPAT from patients in the low PPAT group (Fig. 6B, left panel). Quantification of the signals from F-CHP after 3D reconstruction confirmed the substantial increase in collagen degradation in PPAT from the high PPAT group when compared with the low PPAT group (Fig. 6B, right panel).

**Figure 6:**
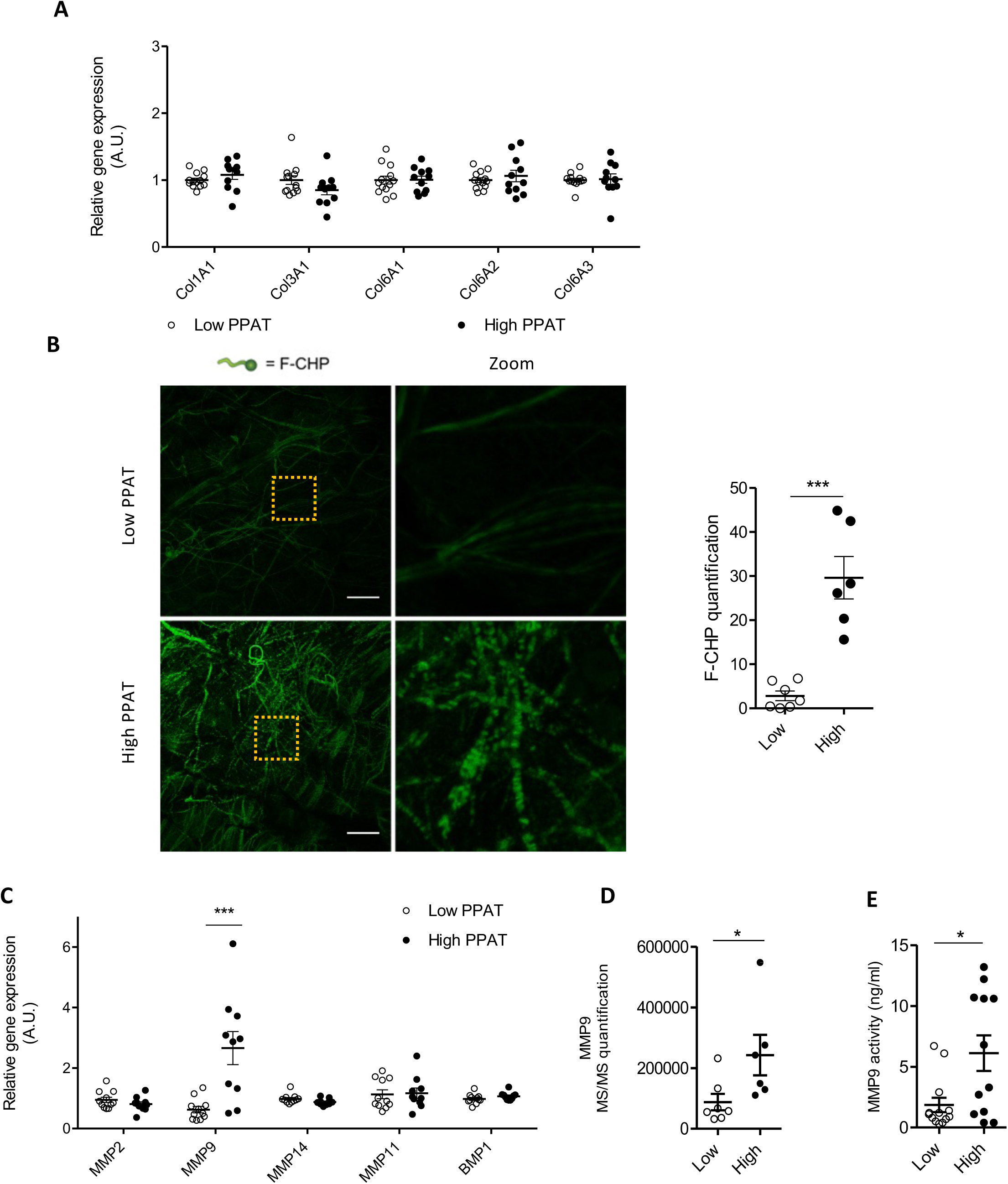
Abundant PPAT exhibits increased collagen degradation likely by MMP9. **A)** Relative expression of genes encoding the indicated collagen isoforms in whole PPAT from patients in the low (n=13) and high qroups (n=11). **B)** Representative confocal images of PPAT stained with fluorescent collagen-hybridizing peptide (F-CHP). The maximum intensity projection of Z-stacked images (left panel) at low magnification and at higher magnification image (Zoom). The areas corresponding to the Zoom images are indicated by the orange dotted lines. Scale bars, 50 μm. F-CHP staining in 3D images was quantified (right panel; n=6-7). **C)** Relative expression of genes encoding the indicated metalloproteinase (MMP) isoforms in PPAT from patients in the low (n=11) and high (n=10) quartiles. **D)** Relative MMP9 protein expression quantified in isolated mature adipocytes by mass spectrometry (n=6-7 per quartile). **E)** MMP9 enzyme activity in protein extracts of PPAT from patients in the low (n=13) and high quartiles (n=12). Bars indicate means ± SEM. Statistical differences were calculated by using Student’s T test. *p < 0.05; *** p< 0.001.

Matrix metalloproteinases are the main enzymes involved in collagen degradation. To investigate the potential involvement of various matrix metalloproteinases in the degradation of collagen in high PPAT, we quantified expression of the main metalloprotease genes expressed in AT (*MMP-2*, *MMP-9*, *MMP-11*, *MMP-14* and *BMP1*) by RT-qPCR. Significantly more *MMP9* mRNA was found in PPAT from patients in the high PPAT group than in PPAT from patients in the low PPAT group (Fig. 6C). This was confirmed by analyzing the proteomics data from isolated adipocytes (Fig. 6D), and it was reflected in elevated MMP9 activity (Fig. 6E). MMP activity is tightly regulated by the balance between expression of the genes encoding MMPs and their endogenous inhibitors the TIMPs (tissue inhibitor of metalloproteinases). Therefore, we analyzed the expression of the genes encoding various TIMP isoforms as well as those encoding the lysyl oxidases LOX and LOXL1, which cross-link ECM proteins. All these genes were expressed at similar levels in the high and low PPAT groups (Fig. S4). We conclude that the smaller amount of collagen in abundant PPAT than in less abundant PPAT and its disorganization is due to enhanced degradation that likely involves MMP9.

### COLVI remodeling is associated with high ETP production in abundant PPAT

Collagen VI is a highly enriched ECM component of AT (28). We investigated whether COLVI was also subjected to degradation in abundant PPATs by immunofluorescence staining and 3D confocal microscopy. As seen in Fig. 7A, the COLVI staining was weaker in the high PPAT group than in the low PPAT group, it was punctiform and the COLVI microfibrils were clearly disorganized, confirming that COLVI is degraded in abundant PPAT. The proteolytic cleavage of the C-terminal part of COL6A3 generates a bioactive fragment called ETP that promotes a wide range of cellular and tissular responses leading to abnormal tissue remodeling, fibrosis and inflammation or tumor progression (19). We then investigated whether ETP generation was increased in abundant PPAT. Tissues were stained with an antibody specific for ETP (30) and imaged by immunofluorescence microscopy (Fig. 7B). Expression of ETP was increased in the high compared to the low PPAT group (Fig. 7B). To investigate the clinical relevance of our findings, we quantified ETP in biological fluids from patients that underwent mpMRI for PCa diagnosis (see Table S4-5 for the description of the new cohort). The patients were classified into the low, mid and high PPAT groups defined above. ETP plasma levels were similar in all patients regardless of their PPAT abundance status (Fig. 7C). By contrast, the ETP concentration in urine collected after prostate massage, which causes exfoliation of cells from the organ into biological fluids (31), was greater in patients in the high PPAT group than it was in patients in the low PPAT group (Fig. 7D). These results confirm the clinical relevance of our findings on ECM remodeling in PPAT from prostatectomy specimens.

**Figure 7:**
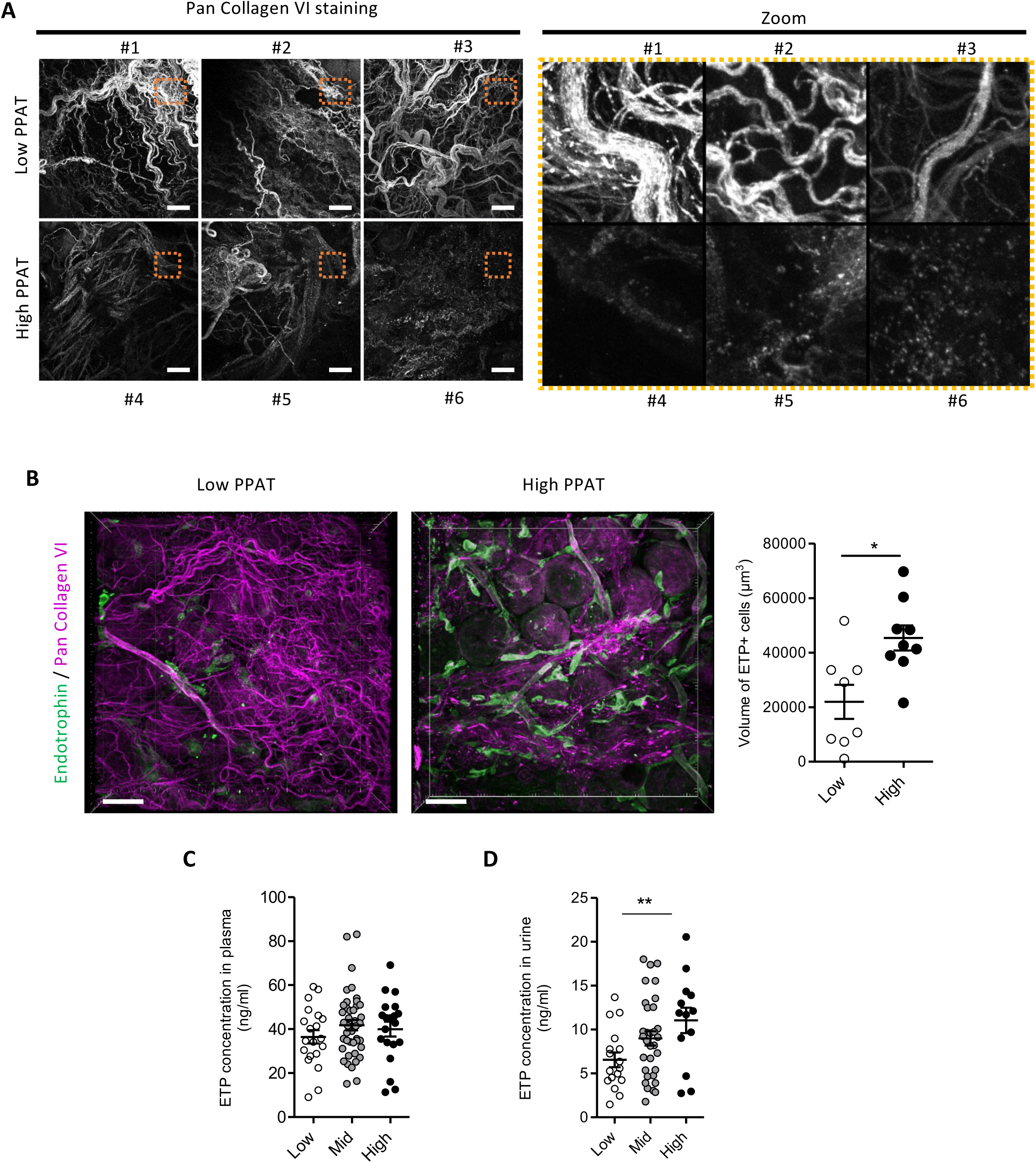
COL6 remodeling is associated with high endotrophin production in abundant PPAT. **A)** Representative confocal images of PPAT from three patients in the low quartile (#1-3) and high quartile (#4-6) stained with an anti-collagen VI antibody that recognizes all collagen VI forms (pan collagen VI). Maximum intensity projections of Z-stacked images at low magnification are shown (left panels) and at higher magnification (Zoom). The areas corresponding to the Zoom images are indicated by the orange dotted lines. Scale bars, 50µm. **B)** Representative confocal immunofluorescence microscopy images of PPAT stained with anti-Pan collagen VI (pink) and anti-endotrophin (green) antibodies. 3D views of Z-stacked images (left). Scale bars, 50µm. Endotrophin staining was quantified on 3D images (right panel; n=8-9). **C)** ETP concentration in plasma from patients with PCa, as quantified by ELISA (Low, n= 20; Mid, n=42; High, n=20). **(D)** ETP concentration in urine from patients with PCa collected after prostate massage, as quantified by ELISA (Low, n= 17; Mid, n=32; High, n=13). Bars indicate means ± SEM. Statistical differences were calculated by using Student’s T test (*p < 0.05 for B) and ANOVA with post-test for linear trend (**p<0.01 for C and D).

## Discussion

One of the most striking features of PPAT is that its accumulation is independent of BMI (6,8). A subset of patients is able to accumulate an excessive amount of PPAT highlighting that PPAT has the capacity to expand upon uncharacterized mechanisms (6,8). These abundant PPATs contribute to the aggravation of prostate-related disorders, especially cancer (6,10–16) despite differences in the methods and criteria that were used to define PPAT abundance. Here, we developed a statistical definition of abundant PPAT as an excess of PPAT volume when compared to the prostate volume predicted from a large cohort of patients. Using this definition, we confirmed that patients with high PPAT exhibit more aggressive PCa tumors in accordance with the literature (6,10–16) and characterized these PPAT at structural and functional levels. Abundant PPATs exhibit extensive ECM remodeling, notably of the collagen network, decreasing the mechanical constraints in hypertrophic adipocytes as shown by the decrease in proteins involved in mechanical sensing and cytoskeleton organization and forces. This expansion in a context of decreased mechanical constraints led to a “stress-free” expansion of the PPATs. As we previously showed that PPAT fibrosis, due to its hypoxic state, limits its expansion, reshaping the ECM is an essential step for its enlargement. This ECM remodeling, with increased expression and activity of the matrix protease MMP9, results in the generation of a bioactive peptide, ETP. The presence of large amounts of ETP in the urine of patients with abundant PPAT reinforces the clinical relevance of our findings. Our main findings and the potential consequences on prostate-related disorders are summarized in Figure 8.

**Figure 8:**
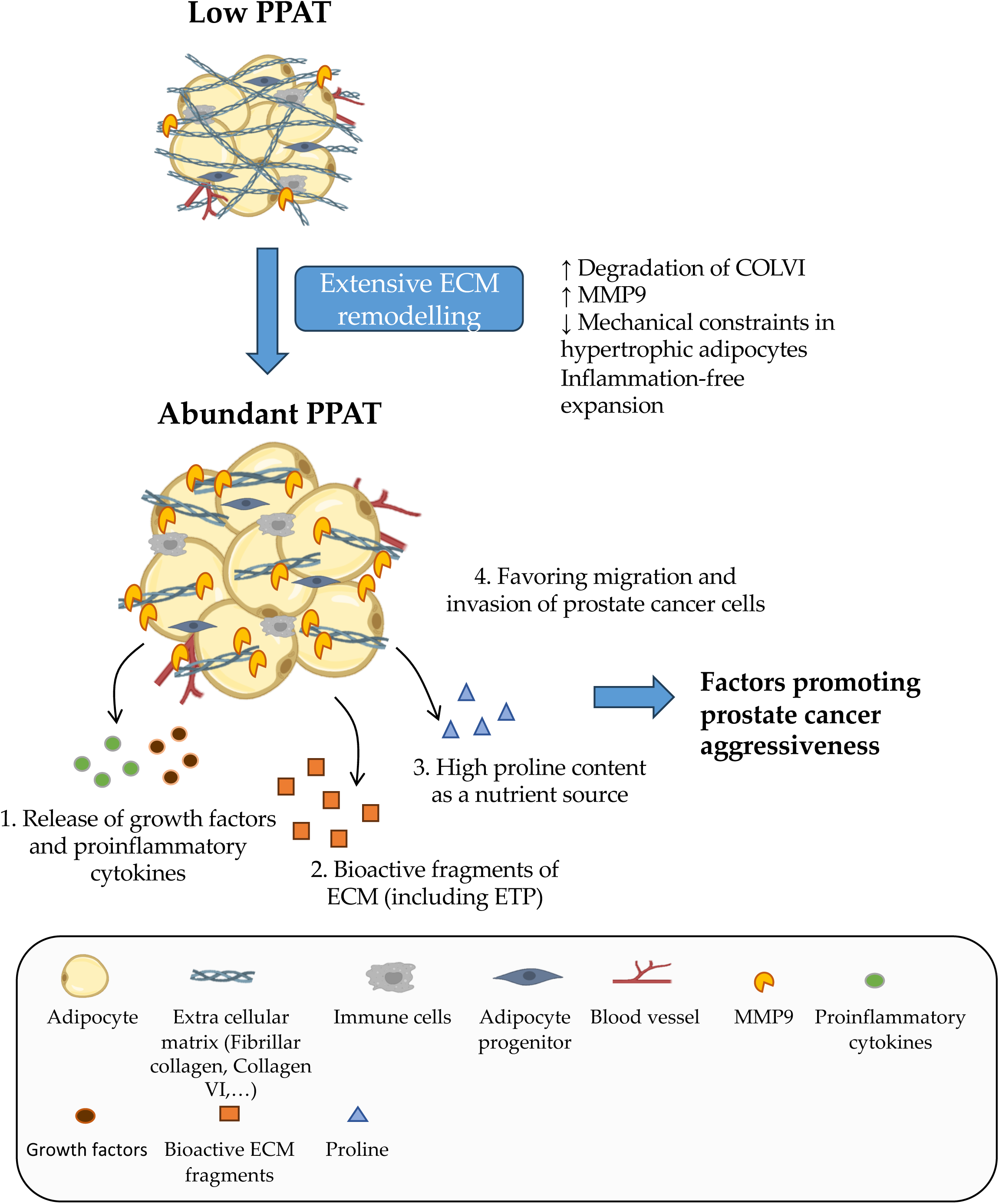
Abundant periprostatic adipose tissue expands through extracellular matrix remodeling resulting in features known to favor PCa progression. As shown before, PPAT exhibits a peculiar vascularization leading to chronic hypoxia and fibrosis that limits its expansion in most individuals. To expand, abundant PPATs exhibit extensive ECM remodeling, notably of the collagen network decreasing the mechanical constraints in hypertrophic adipocytes leading to an inflammation free-expansion. The degradation of ECM and particularly COL VI, likely by MMP-9, contributes to the overexpression of a bioactive fragment, endotrophin, as shown *ex vivo* and in the urine of patients with abundant PPAT. This unique mechanism of PPAT expansion results in features known to favor prostate-related disorders including PCa. The degradation of the ECM could release growth factors, pro-inflammatory cytokines (1), bioactive ECM fragments (including ETP) (2) and/or increase the availability of proline which could act as a nutrient source for cancer cells (3). Finally, the loose collagen network could promote prostate cancer cells’ migratory and invasive capacities (4).

In this work, we first developed a new approach to define PPAT abundance using mpMRI – the gold standard for imaging the prostate and surrounding tissues – to measure both prostate volume and the whole volume of PPAT rather than simply the thickness of fat or the area of the tissue at a single point. We measured these volumes by applying a slice-by-slice segmentation technique that we used in a previous study (8). In accordance with previous studies (17,18) we demonstrate that the volume of the PPAT correlates with that of the prostate. As prostate size is highly variable between individuals and often increases with age (32), it is important to take this parameter into account to accurately define the relative fat volume in the periprostatic area. Recent studies have used NPFV to compare PPAT abundance between individuals in order to better define the relative fat volume in the periprostatic area (14,15,17,18). However, this approach might introduce a bias towards patients with smaller prostates being identified as having abundant PPAT, as found in a previous study (17). To overcome this bias, we developed a linear regression model that normalizes PPAT abundance by comparing the residual value of PPAT volume to the linear relationship between PPAT and prostate volume computed from the whole cohort. Using this approach, we calculated PPAT abundance for each individual in the cohort, ranked the population according to PPAT abundance and divided it into quartiles. We observed no differences in prostate size between the patients in these quartiles, thus validating our approach. Applying our new approach to define PPAT abundance, we found that patients with the most abundant PPAT (high PPAT group), have the most aggressive PCa (as defined by ISUP scores and percentage of high-grade lesions). Therefore, our new and robust definition of PPAT abundance can be applied to future translational studies on prostate-related disorders including PCa. Using this validated definition of PPAT abundance, we investigated whether PPAT characteristics from the different groups differ in their histology, transcriptomes and proteomes in order to understand how abundant PPATs expand.

The discovery that abundant PPAT contains a loose meshwork of both fibrillar and non-fibrillar collagens is perhaps the most important finding from this study because it explains how the tissue expands and why this expansion occurs without any further increase in inflammation nor with the changes in adipocyte secretions that are usually observed in dysfunctional AT. We previously demonstrated that PPAT exhibits fibrosis preventing its expansion in obese patients and therefore reshaping the ECM is an essential step for its enlargement (8). In fact, fibrosis, by generating a mechanical stress on adipocytes, plays a central role in limiting the expandability of AT (28). One of the main differences we saw between the ECM in the high and low PPAT groups was in COLVI, the most abundant non-fibrillar collagen in AT (28). Whereas COLVI staining revealed a typical beaded filament network (33) in the PPAT of the low PPAT group, in the high PPAT group the staining was weak and punctiform and the filament network was disorganized. COLVI interacts with many ECM proteins, including fibronectin and collagen types I, II and IV, bridging cells to the surrounding connective tissue and organizing the three-dimensional tissue architecture (33). Thus, the overall loss of collagen may result from initial extensive degradation of COLVI. Degradation of COLVI leads to the overexpression of ETP, that was found to be higher in the urine of patients with abundant PPAT confirming the clinical relevance of our results. COLVI degradation probably involves MMP9, which we found was much more abundant in the high PPAT group than in the low PPAT group. Degradation of COLVI by MMP9 has been described previously (34), as well as by MMP2 (34), MMP-11 (35) and MMP14 (36). Expression of the latter MMPs and the inhibitors TIMP 1-4 were similar in both groups, however, suggesting that MMP9 is the most important in degrading COLVI in abundant PPAT. In addition, this is consistent with a report showing that *in vitro* MMP9 activity, along with MMP2 and MMP16, is key to produce active ETP (37). We do not know yet what initiates ECM remodeling to allow PPAT to expand. The correlation between the volume of PPAT and the volume of the prostate suggests that steroid hormones might be involved. Consistent with this, MMP9 expression is regulated by androgens (38) and **s**ex hormones influence the metabolism, endocrine functions and angiogenesis of certain other ATs (39), so this hypothesis deserves further investigation.

The loose collagen network, we saw in abundant PPAT, will be permissive to adipocyte to enlarge and in fact we find that abundant PPATs exhibit adipocyte hypertrophy. This hypertrophy may be explained by both the increased expression of lipid transporters involved in uptake of lipids into adipocytes and the decreased expression of proteins involved in lipolysis, favoring lipid uptake over degradation. The expansion of PPAT we observed in PCa patients in our high PPAT group does not result in tissue inflammation or in adipocyte dysfunction. We showed that the loose meshwork of fibrillar and non-fibrillar collagens in high PPAT, resulting from enhanced ECM degradation, decreases the expression in adipocytes of proteins involved in mechano-transduction and stiffness sensing. This expansion in a context of decreased mechanical constraints led to a “stress-free” expansion allowing the adipocytes to enlarge without cell death, which prevents accumulation of macrophages and, subsequently, the occurrence of an inflammatory microenvironment. Evidence from mouse models deficient in ECM or overexpressing ECM remodeling proteins, support our interpretation. Mice that lack COLVI and that are obese due to either a high fat diet (HFD) or to *ob/ob* mutation have impaired ECM stability and, hence, reduced AT fibrosis resulting in adipocyte hypertrophy in the absence of inflammation (42). Similarly, in an inducible AT specific MMP14 overexpression model, the overexpression of MMP14 during the early stage of HFD reduced inflammation and fibrosis. In both cases, this phenotype was linked to the decrease in ECM stiffness of AT, thereby releasing the mechanical stress to allow for its healthy expansion (42,43). Taken together, our results demonstrated that PPAT expands by a unique mechanism that protects it from further dysfunction.

The extensive ECM remodeling we observed in abundant PPAT results in features known to favor PCa and other prostate-related disorders (including benign prostate hyperplasia (BPH), erectile and urethral dysfunctions). First, ECM functions as a reservoir by binding numerous growth factors and pro-inflammatory cytokines that are released by its remodeling susceptible to act on adjacent normal and cancer prostate cells (44). Second, components of the ECM also interact with epithelial cells by serving as ligands for cell receptors such as integrins, transmitting signals that regulate adhesion, migration, proliferation, apoptosis, survival or differentiation (44). Among the components produced by enzymatic cleavage of ECM proteins, we identified a higher expression of ETP, a bioactive fragment derived from proteolytic cleavage of the carboxy-terminal C5 domain of the COLVI α3 chain from the parental COLVI microfibrils during secretion (45). Adipocyte-derived ETP favors breast tumor progression and resistance to therapy in genetically engineered mice (45) and in xenografted human tumors (30). Third, degradation of collagen might contribute to act as a proline reservoir for proximal epithelial cells and serve as a nutrient source (46). Finally, structural reorganization of the ECM observed in abundant PPAT might favor the migratory capacities of adjacent PCa cells (47).

Deciphering these mechanisms require a model capable of reproducing the interaction between abundant PPATs and normal or tumoral epithelial cells. Coculture of primary PPAT explants with PCa cells might represent an attractive approach but explant culture is associated with rapid increase in pro-inflammatory cytokines and substantial dedifferentiation of adipocyte resulting in altered pattern of adipocyte marker gene expression as previously described (48) and as we observed for PPAT (data not shown) rendering this approach unsuitable. Alternative models have been proposed such as the so-called “sandwiched AT” where human AT fragment is enclosed between two monolayers of adipose stromal cells ensuring their viability, metabolic and secretory function for up to 8 weeks (49). Setting up this approach on PPAT could be interesting although time consuming.

In conclusion, we have deciphered for the first time the original mechanism that allows human PPAT to expand and demonstrated that it relies on ECM reshaping that is, to our knowledge, a unique mechanism of expansion among adipose depots. This work opens new avenues to decipher the role of abundant PPAT in PCa at both fundamental and clinical levels. Since mpMRI is now widely used prior to biopsies, dedicated clinical trials should be designed to assess the interest of PPAT abundance in risk stratification based on, in addition to its volume, dedicated radiomic studies to corroborate to ECM content as well as by the search for urinary surrogate markers linked to ECM degradation.

## Materials and Methods

### Patients, prostate and fat measurements

Between September 2016 and July 2021, patients with localized PCa diagnosed by prostate biopsies who were opting for radical prostatectomy (RP) were recruited. We excluded from the study patients with metastatic PCa (as assessed by computerized tomography and/or bone scan), and patients who had received radiotherapy, hormonal or high intensity ultrasound treatment. All the patients underwent pre-operative 1.5 T mpMRI after injection with 20 mg of butyl scopolamine (Buscopan; Boehringer-Ingelheim, Paris, France). Anatomic 3D fast spin echo T2-weighted MRI, functional diffusion-weighted MRI and dynamic contrast-enhanced MRI data were acquired. After anonymization of the mpMRI data obtained from the medical records of the patients, the PPAT and prostate volume, the subcutaneous and perirectal AT area was measured using a semi-automated segmentation technique on contiguous 3 mm T2-weighted axial slices and Olea Sphere software (Olea medical, La Ciotat, France). The subcutaneous AT area was determined by measuring the perpendicular distance between the skin and the anterior upper border of the symphysis pubis on a selected T2-weighted axial slice where it was maximum. The perirectal AT area was determined by segmenting the fat around the rectum at the level of the apex of the prostate. (Table S1). Twenty patients who underwent MRI for suspected PCa with negative results were included in this radiological study for the measure of PPAT and prostate volume (named patients no cancer). Three observers, trained by a senior experienced radiologist and blind to the clinical and pathological data, carried out all segmentations and measurements. The PPAT was segmented from the level of the prostate base to the apex (see Figure 1A) (mean 14 slices per patient) as previously described (8). Baseline demographic characteristics (age and BMI), serum Prostate Specific Antigen (PSA) concentration, pathological parameters of the tumor on RP pieces including the volume of the tumor, Gleason score and ISUP group (histological score reflecting the differentiation of PCa determined by a trained pathologist according the 2016 WHO classification and ISUP recommendations), percentage of high grade lesions (defined as the percentage of the tumor with a Gleason score greater than 4), pT (pathological T) stage and extraprostatic extension were acquired from the medical records of the patients. The clinical characteristics of the cohort is presented in Table S2. The Gleason score and ISUP group was also determined on pre-operative biopsies by a trained pathologist.

### PPAT collection and tissue processing

PPAT from patients undergoing RP were removed and the samples were anonymized. Charred samples and those weighing < 300 mg were excluded from the study. Samples were immediately put in 50 mL tubes containing 10 mL of Dulbecco’s Modified Eagle’s Medium (DMEM, Thermofischer Scientific, Courtaboeuf, France) and were transported to the research lab within 1 h. The samples were weighed and use for several preparations. For RT-qPCR analysis, measure of MMP9 activity and dosage of hydroxyproline content, PPAT were immediately frozen in liquid nitrogen and stored at – 80°C for later use. For 3D microscopy, samples were fixed with 4% paraformaldehyde solution at room temperature (RT) for 24 h. Some of the samples were also used to isolate adipocytes and the stromal vascular fraction after collagenase digestion (see below). Finally, samples were used to prepare PPAT-conditioned medium as previously described

### Isolation of AT stromal vascular fraction (SVF) and mature adipocytes

Digestion of fresh PPAT was performed as previously described (8). Briefly, PPAT were digested with 250 U/ml type I collagenase (Sigma-Aldrich, Saint-Quentin-Fallavier, France) in PBS containing 2 % BSA for 30 min at 37°C under constant shaking. The samples were filtered with a 250 µm cell strainers and the cell suspensions were then gently centrifuged for 1 min at 20 g at RT to pack adipocytes on the top layer. Floating adipocytes were collected and rinsed with KRB buffer supplemented with 1mM Hepes and 0.5% FFA free-BSA (all obtained from Sigma-Aldrich, Saint-Quentin-Fallavier, France) to obtain pure adipocyte cell suspensions that were used for proteomic analysis (see below). After removal of adipocytes, the remaining cell suspension was centrifuged at 300 g for 10 min to pellet SVF cells. Erythrocytes were lysed by incubation at RT in erythrocyte lysis buffer (155 mM NH4Cl, 5.7 mM K2HPO4, 0.1 mM EDTA, pH 7.3) for 10 min followed by successive filtration through 100, 70, and 40 µm strainers. The viable recovered cells were washed 2 times in PBS, counted and analyzed by flow cytometry.

### Flow cytometry analysis

Flow cytometry was performed as previously described with minor modifications (22). Briefly, 1 x 10^5^ stromal vascular cells were incubated for 30 min at 4 °C in PBS supplemented with 0.5% BSA and 2 mM EDTA containing fluorescently labeled antibodies. Brilliant Violet 510™ anti-human CD45 (clone HI30, dilution 1/20, Biolegend, Amsterdam, The Netherlands), APC anti-human CD206 (clone 19.2, dilution 1/10, BD Biosciences, Le Pont de Claix, France) and PE-Vio® 770 anti-human CD14 (clone TÜK4, dilution 1/50, Miltenyi Biotec, Paris, France) antibodies were used for macrophages phenotyping. PerCP anti-human CD34 (Clone 8G12, dilution 1/20, BD Biosciences, Le Pont de Claix, France), BV450 anti-human CD31 (Clone WM59, dilution 1/20, BD Biosciences, Le Pont de Claix, France), PE anti-human MSCA1 (Clone W8B2, dilution 1/10, Miltenyi Biotec, Paris, France), APC anti-human CD271 (Clone ME20.4-1.H4, dilution 1/10, Miltenyi Biotec, Paris, France) and Brilliant Violet 510™ anti-human CD45 (clone HI30, dilution 1/20, Biolegend, Amsterdam, The Netherlands) antibodies were used for progenitor cell subset phenotyping. Brilliant Violet 510™ anti-human CD45 (clone HI30, dilution 1/20, Biolegend, Amsterdam, the Netherlands), Brilliant Violet 450 anti-human CD3 (Clone UCHT1, dilution 1/20, BD Biosciences, Le Pont de Claix, France), FITC anti-human CD4 (Clone RPA-T4, dilution 1/20, BD Biosciences, Le Pont de Claix, France), PerCP anti-human CD8 (Clone SK1, dilution 1/10, BD Biosciences, Le Pont de Claix, France) and APC anti-human CD25 (Clone 2A3, dilution 1/10, BD Biosciences, Le Pont de Claix, France) antibodies were used for lymphocyte phenotyping. As control, a panel of appropriate isotype antibodies was used. The labeled cells were washed with PBS and analyzed in a FACS Canto^TM^ II flow cytometer using Diva Pro software (BD Biosciences, Le Pont de Claix, France). Cell debris, dead cells and doublets were excluded based on scatter signals. Compensation was established using compensation particles set (BD Biosciences, Le Pont de Claix, France) and single staining and flow cytometer calibration was performed with rainbow calibration particles (BD Biosciences, Le Pont de Claix, France).

### ELISA assays

As previously described (8), a panel of secreted adipokines were quantified in PPAT-CM with an ELISA kit (LXSAHM-08; R&D Systems, Noyal Châtillon sur Seiche, France) and a mixture of antibodies against Adiponectin/Acrp30, IL-6, IL-1β, Leptin, MCP1 and TNFα, according to the protocol provided by the manufacturer. Endotrophin was quantified in plasma and urine collected after prostate massage using homemade ELISA as previously described (30).

### RNA extraction and RT-qPCR

For RNA extraction, 100 mg of frozen tissue was mixed with 500 µl QIAzol^TM^ lysis reagent (Qiagen, Courtaboeuf, France) and homogenized with a Precellys ® tissue homogenizer (2 cycles of 45 seconds at 7500 rpm at 4°C) using the Cryolys cooling system. The tissue homogenates were centrifuged for 1 min at 5000 g at RT. The lipid-containing surface layer was then removed and 350µl of chloroform was added. The organic and aqueous phases were separated by centrifugation (10 min at 5000 g at RT). The upper phase, containing RNA, was recovered and mixed with ethanol for RNA extraction using the Qiagen RNeasy mini kit as described by the manufacturer (Qiagen, Courtaboeuf, France). The RNA concentration was determined with a Nanodrop 2000 spectrophotometer at a wavelength of 260nm. cDNA were reverse transcribed from 250 ng of purified RNA by using the RevertAid H Minus First Strand cDNA synthesis kit (Thermo Fisher Scientific, Illkirch-Graffenstaden, France) with hexamer random primers according to the supplier’s recommendations. Gene expression was quantified by RT-qPCR on 6.25 ng of cDNA with TaqMan primers (the primers used are listed in Table S6), Thermo Fisher Scientific, Illkirch-Graffenstaden, France) and Fast Advanced TaqMan Master Mix (Thermo Fisher Scientific, Illkirch-Graffenstaden, France) in a final volume of 20 µl with a CFX96real time PCR system (Bio-Rad). Results were normalized to the amount of 18S rRNA.

### Sample preparation for proteomic analysis

Isolated adipocytes obtained as described above were washed in PBS three times and proteins were extracted from 500 μl of adipocyte with 500 μl 1X RIPA buffer (Sigma-Aldrich) supplemented with complete antiprotease tablet (Roche) with a precellys tissue homogenizer and Ceramic beads (CK14) with 2 cycles of 45 s at 7500rpm at 4°C. The samples were then centrifuged 10 min at 10 000 g at RT to separate and aggregate the lipids at the top layer from which they were removed. The extract was recovered without the top layer containing the lipids and centrifuged again 10 min at 10 000 g at 4°C. Protein concentration was determined using DC protein assay kit (Biorad, Biorad, Marnes-la-Coquette, France) following manufacturer instructions. 40μg of proteins was digested for proteomic characterization by mass spectrometry using the S-Trap™ micro spin column (ProtiFi, NY USA).

### LC-MS/MS analysis

Tryptic peptides were resuspended in 40 µl of 2% acetonitrile and 0.05% trifluoroacetic acid and analyzed by nano-liquid chromatography using an UltiMate 3000 system (NCS-3500RS Nano/Cap System; Thermo Fisher Scientific) coupled to an Orbitrap Q Exactive HF-X mass spectrometer (Thermo Fisher Scientific). Peptides were separated on an Acclaim PepMap C18 column (id 75 µm, length 50 cm, particle size 2 µm, Thermo Fisher Scientific) using a 10%-45% gradient of solvent B (80% acetonitrile, 0.2% formic acid), over 120 min (solvent A: 5% acetonitrile, 0.2% formic acid). The mass spectrometer was operated in data-dependent acquisition mode with the Xcalibur software.

### Bioinformatic MS data analysis

Acquired MS and MS/MS data were searched with Mascot (version 2.6.1, http://matrixscience.com) against the Human UniProtKB database (Swiss-Prot/TrEmbl, 556,568 entries). Cysteine carbamidomethylation was set as a fixed modification. Methionine oxidation and acetylation of protein N-terminus were set as variable modification. Up to two missed trypsin/P cleavages were allowed. Mass tolerances in MS and MS/MS were set to 10 ppm and 0.8 Da, respectively. Proline software (version 1.6, http://proline.profiproteomics.fr/) was used for the validation of identifications and the label-free quantification of identified proteins (50). Mascot identification results were imported into Proline. Search results were validated through a false-discovery rate (FDR) set to 1% at protein and peptide-sequence match level. Proteins with a p-value < 0.05 and a ratio of average normalized area < 0.5 and > 2 were considered significant. Volcano plots were drawn to visualize significant protein abundance variations between two conditions control and assay. They represent log10 (p-value) according to the log2 ratio. Label free quantification intensities were normalized between samples with the function “NormalizeBetweenArrays” from limma package (v3.40.6) in R (v3.6). Differential analysis was computed with limma package from Bioconductor. Principal component analysis was performed with R (v3.6) and FactoMineR package (v2.4). Go term enrichment analysis was performed with Molecular signature database on differentially represented proteins (V7.4 MSigDB).

### 3D confocal microscopy and quantification

For immunofluorescence staining, PPAT (250mg) were saturated and permeabilized with PBS supplemented with 3 % BSA and 0.2 % Triton X100 (Sigma-Aldrich, Saint-Quentin-Fallavier, France) for 1 h. Tissues were then incubated overnight at 4°C with the following primary antibodies diluted in PBS 3% BSA 0.2% Triton X100: anti-β-Tubulin (clone 2A1A9, dilution 1/200, Abcam, Cambridge, UK), anti-YAP1 (clone D8H1X, dilution 1/200, Cell Signaling Technology, Leiden, The Netherlands), Anti-Pan Collagen VI (clone 3C4, dilution 1/200, Millipore, Guyancourt, France) and serum anti-human ETP (30) (dilution 1/100) antibodies. Tissues were washed 5 times with PBS – 0.05% Tween for 10 min. Then, tissues were incubated with secondary antibodies coupled with fluorescent dye (dilution 1/800, Biotium, Fremont, CA, USA) according to the species in which the primary antibodies were produced for 3h at RT and with or without rhodamine-phalloidin (dilution 1/200, Thermofisher, Waltham, MA, USA) diluted in PBS, 3% BSA and 0.2% Triton X100. Tissues were washed 5 times with PBS 0.05% Tween for 10 min. Nuclei were stained with Hoechst 33342 (dilution 1/1000, Invitrogen, Villebon sur Yvette, France) for 30 min and after 3 washes in PBS Z stack images were acquired using LSM710 confocal microscope and 40X objective. For Fluorescent Collagen Hydridizing Peptides (F-CHP, Cat@FLU-300, Clinisciences, Nanterre,France), fixed AT were incubated with 20µM of F-CHP overnight at 4° C. Tissue were rinsed with PBS 5 times for 5-10 min. Z stack images were acquired using LSM710 confocal microscope and 40X objective. Staining was quantified with Imaris software (v9.2). Briefly, after segmentation and thresholding of the staining of interest, volumes were rendered with the surface module of Imaris. Extracted statistics were normalized to the volume of the tissue determined with the autofluorescence of the tissue.

For Picrosirius Red, fixed AT were incubated for 3 h in Picrosirius Red solution (1 g/L Sirius red in 1.3% picric acid). After two washes in acidified water followed by fixation and dehydration steps in 100% ethanol, images were acquired with a confocal microscope (LSM710, Zeiss International, Oberkochen, Germany) equipped with a 40 oil-immersion inverted objective with a numerical aperture of 1.3. The samples were excited at 560 nm, and fluorescence emission was collected above 580 nm. Pinhole was setup at 0.7 Airy (ZEN imaging software; Zeiss International) to increase resolution. Three-dimensional reconstructions and analyses of collagen fiber organization were performed with Imaris software and the filament tracer module. The data generated were normalized to the tissue volume obtained after tissue surface reconstruction with Imaris (v9.6) (Bitplane, Gometz la Ville, France).

### Hydroxyproline quantification

Fragments of PPAT of about 100 mg were weighed precisely, placed in 500 µl deionized water in a Precellys® 2mL Soft Tissue homogenizing Ceramic beads Kit (CK14) (Ozyme, Saint-Cyr-l’École, France) and extracted by using the Precellys® tissue homogenizer with 2 cycles of 45 seconds at 7500 rpm at RT. Samples were then centrifuged for 10 min at 5000 g at RT to remove lipids floating on the top layer by pipetting. Lipid-free supernatants were vortex mixed, 150 µl were mixed with an equal volume of 10 M NaOH and heated at 105°C for 1 hour to fully digest collagens. The reaction was stopped by adding 150 µl of 10M HCl. Hydroxyproline was quantified in duplicate by using the Hydroxyproline Assay Kit (Perchlorate-Free; Milpitas, CA, USA) as recommended by the supplier and normalized to the precise tissue weight initially digested.

### MMP9 activity

For protein extraction, 100-250 mg of frozen tissues were incubated with 400 µl of RIPA lysis buffer (Sigma-Aldrich, Saint-Quentin-Fallavier, France). Proteins were extracted using the Precellys ® tissue homogenizer and Soft Tissue homogenizing Ceramic beads (CK14) with 2 cycles of 45 seconds at 7500 rpm at RT. The samples were then centrifuged 10 min at 5000 g at RT to separate and aggregate the lipids on the top layer. The lipid-free phase was recovered, sonicated and centrifuged again 10 min at 10 000 g at 4°C. Protein concentration was quantified with DC protein assays (Bio Rad, Marnes-la-Coquette, France). Concentration of active MMP9 on 200µg of total proteins lysate was quantified by using a Human Active MMP-9 Fluorokine E assay kit (F9M00; R&D Systems, Noyal Châtillon sur Seiche, France) according to the protocol provided by the manufacturer. Fluorescence was quantified using FLX800 fluorimeter with an excitation wavelength set to 340nm and an emission wavelength set to 405nm.

## Statistics

Univariate statistical analysis of clinical data was performed using ANOVA for continuous variables or using Chi square test for qualitative variables with Prism (v5.04). Linear regression was computed with Prism v5.04 and residuals were calculated according to the fitted linear model. Multivariate analysis was performed by multiple logistic regression using the glm function from R (v3.6) to predict patients having an ISUP score above or equal to 3. For experimental data, statistical analysis was performed using Student T test after testing normality distribution using Kolgomorov-Smirnov test for comparison between two groups or with ANOVA followed by test for linear trends between 3 groups. Statistical differences were considered as significant if the p-value was below 0.05.

## Study approval

The study was conducted in accordance with the guidelines and with the full approval of the national ethics committee (AC-2020-4031). Written informed consent was received from participants before inclusion in the study, which was conducted in accordance with the Declaration of Helsinki principles as revised in 2000.

## Data availability

The datasets used and analyzed during the current study are available from the corresponding author on reasonable request. The mass spectrometry proteomics data have been deposited to the ProteomeXchange Consortium via the PRIDE partner repository with the dataset identifier PXD038514. Reviewer account details: Username; Password: 5QSTgkLK.

## Author contribution

DE and AT conducted and analyzed most of the experiments presented in the manuscript and prepared the figures. AT, MR and DM performed the segmentations and measurements of PPAT and prostate volumes. DE developed the statistic model defining the PPAT abundance. CF performed the H&E staining of PPAT samples and AT performed the adipocyte size measurement. AT performed the flow cytometry experiments with the help of CB under the supervision of AB. AT and MH performed the ELISA experiments with the help of SLG under the supervision of PV. DE performed the proteomic experiments and MDP performed the analysis of the proteomic data under the supervision of OBS. CH and YJ performed the qPCR experiments and MMP9 activity measurement. DE performed all the microscopy experiments with the help of SP, NVA and SD. DB performed the endotrophin dosage in human fluids under the supervision of PES. ML helped to write the manuscript and to prepare the figures. BM, CM, ND, MT, MR harvested the human periprostatic tissue and fluids samples and CM and MR collected the clinical data. MM participated to tissue processing. DE, DM and Catherine Muller (CM) supervised the study and wrote the manuscript. All the authors reviewed the manuscript. All the authors agreed with the final version of the manuscript.

## Acknowledgements

This work benefited from the Toulouse Réseau Imagerie (TRI)-RIO Optical Imaging Platform at the Institute of Pharmacology and Structural Biology (Genotoul, Toulouse, France). We acknowledge Life Science Editors for professional scientific editing during the preparation of the manuscript.

This study was supported by the Ligue Nationale Contre le Cancer (Équipe Labélisée), the French National Cancer Institute (INCA PL-BIO 2016-176 to AB, BM, PV, and CM), by the Fondation Toulouse Cancer Santé (to AB, BM, OBS, PV and CM) and by the Fondation ARC (Association de Recherche contre le Cancer) Programmes Labelisés 2016 (to AB, PV and CM). The work was also funded in part by the French Ministry of Research with the Investissement d’Avenir Infrastructures Nationales en Biologie et Santé program (ProFI, Proteomics French Infrastructure project, ANR-10-INBS-08) (to OBS). DE received a post-doctoral fellowship from the Fondation pour La Recherche Médicale (SPF201809007124).

---

**Table S1:** Anthropometrical characteristics of the PCa patients.

| Characteristic | N = 351<sup>1</sup> |
| --- | --- |
| Age (years) | 65 (60, 69) |
| BMI (kg/m<sup>2</sup>) | 25.48 (24.21, 27.76) |
| *Missing* | 19 |
| PPAT volume (cm<sup>3</sup>) | 43 (34, 53) |
| Subcutaneous AT area (cm<sup>2</sup>) | 35 (28, 44) |
| *Missing* | 156 |
| Prostate volume (cm<sup>3</sup>) | 44 (35, 60) |
| Perirectal AT area (cm<sup>2</sup>) | 25.9 (22.4, 30.3) |
| *Missing* | 96 |

<sup>1</sup>Data are presented as the median value with inter-quartile range (IQR) in brackets. BMI, Body mass index

**Table S2:**
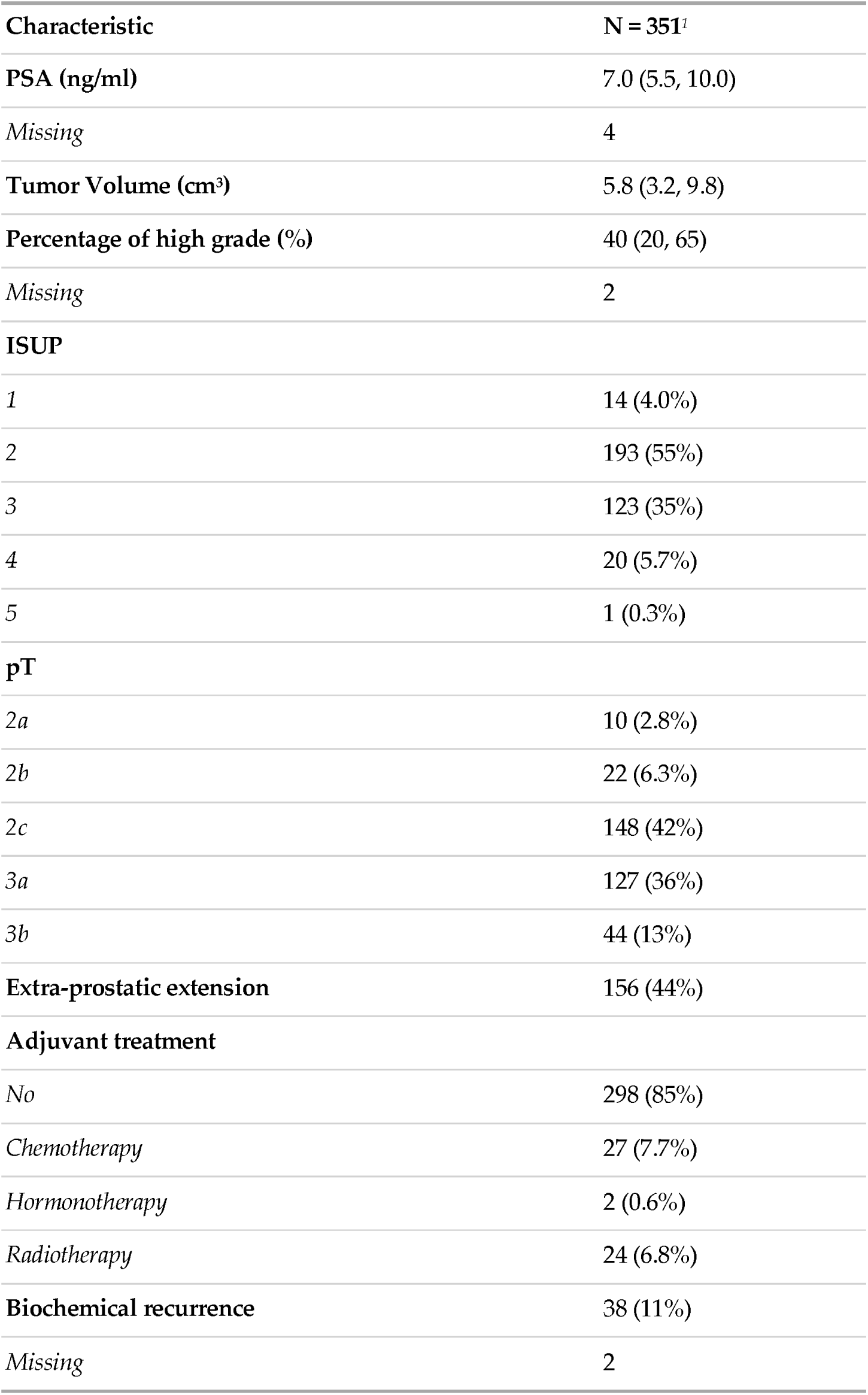
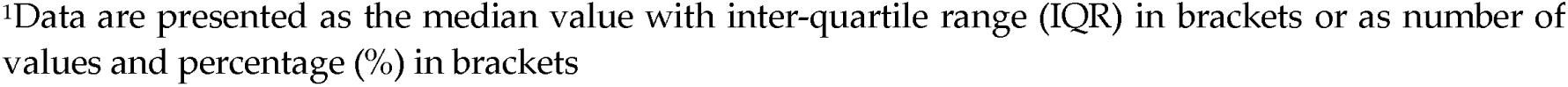
Clinical and pathological characteristics of the PCa patients.

| **Characteristic** | **N = 351<sup>1</sup>** |
| --- | --- |
| **PSA (ng/ml)** | 7.0 (5.5, 10.0) |
| *Missing* | 4 |
| **Tumor Volume (cm<sup>3</sup>)** | 5.8 (3.2, 9.8) |
| **Percentage of high grade (%)** | 40 (20, 65) |
| *Missing* | 2 |
| **ISUP** |  |
| 1 | 14 (4.0%) |
| 2 | 193 (55%) |
| 3 | 123 (35%) |
| 4 | 20 (5.7%) |
| 5 | 1 (0.3%) |
| **pT** |  |
| 2a | 10 (2.8%) |
| 2b | 22 (6.3%) |
| 2c | 148 (42%) |
| 3a | 127 (36%) |
| 3b | 44 (13%) |
| **Extra-prostatic extension** | 156 (44%) |
| **Adjuvant treatment** |  |
| *No* | 298 (85%) |
| *Chemotherapy* | 27 (7.7%) |
| *Hormonotherapy* | 2 (0.6%) |
| *Radiotherapy* | 24 (6.8%) |
| **Biochemical recurrence** | 38 (11%) |
| *Missing* | 2 |

<sup>1</sup>Data are presented as the median value with inter-quartile range (IQR) in brackets or as number of values and percentage (%) in brackets

**Table S3:** Clinical and pathological characteristics of the PCa patients according to PPAT abundance.

| Characteristic<sup>1</sup> | Low, N = 88 | Mid, N = 176 | High, N = 87 | p-value |
| --- | --- | --- | --- | --- |
| PSA (ng/ml), Median (IQR) | 7.0 (5.1, 9.6) | 7.0 (5.2, 9.0) | 8.0 (6.1, 11.1) | 0.0179<sup>2</sup> |
| (Missing) | 1 | 3 | 0 |  |
| Tumor Volume (cm<sup>3</sup>), Median (IQR) | 5.1 (2.8, 9.5) | 6.2 (3.3, 9.9) | 6.0 (3.0, 9.9) | 0.4400<sup>2</sup> |
| Percentage of high grade (%), Median (IQR) | 28 (10, 60) | 40 (20, 70) | 55 (22, 72) | 0.0008<sup>2</sup> |
| (Missing) | 0 | 2 | 0 |  |
| ISUP ≥ 3, n (%) | 26 (30) | 73 (41) | 45 (52) | 0.0115<sup>3</sup> |
| Missing | 21 | 56 | 30 |  |
| pT n (%) |  |  |  | 0.2413<sup>3</sup> |
| 2a or 2b | 6 (6.8) | 17 (9.7) | 9 (10) |  |
| 2c | 44 (48) | 67 (38) | 39 (45) |  |
| 3a | 35 (40) | 66 (38) | 26 (30) |  |

PSA, prostate specific antigen

|  |  |  |  |  |
| --- | --- | --- | --- | --- |
| *3b* | 5 (5.7) | 26 (15) | 13 (15) |  |
| **Extra-prostatic extension** | 39 (44) | 81 (46) | 36 (41) | 0.7752<sup>3</sup> |

<sup>1</sup>Data are presented either as the median value with inter-quartile range (IQR) in brackets or as number of values and percentage (%) in brackets. PSA, prostate specific antigen
Kruskal-Wallis rank sum test
<sup>3</sup> Pearson's Chi-squared test

**Table S4:** Anthropometrical characteristics of the patients in the cohort used for endotrophin quantification in biological fluids.

| Characteristic | N = 78<sup>1</sup> |
| --- | --- |
| Age (years) | 66.0 (61.8, 69.0) |
| *Missing* | 6 |
| BMI (kg/m<sup>2</sup>) | 25.68 (24.32, 27.15) |
| *Missing* | 12 |
| PPAT Volume (cm<sup>3</sup>) | 41 (33, 52) |
| Prostate Volume (cm<sup>3</sup>) | 48 (38, 62) |
| PPAT Abundance (Group) |  |
| *Low* | 20 (26%) |
| *Mid* | 37 (47%) |
| *High* | 21 (27%) |
| <sup>1</sup> |  |

<sup>1</sup>Data are presented as the median value with inter-quartile range (IQR) in brackets. BMI, Body mass index

**Table S5:** Clinical and pathological characteristics of the patients in the cohort used for endotrophin quantification in biological fluid.

| Characteristic | N = 78<sup>1</sup> |
| --- | --- |
| **PSA (ng/ml)** | 6.0 (4.0, 10.0) |
| *Missing* | 1 |
| **Tumor Volume (cm<sup>2</sup>)** | 3 (2, 6) |
| *Missing* | 37 |
| **Undifferentiated cancer** | 30 (10, 65) |
| *Missing* | 6 |
| **ISUP** |  |
| 0 | 1 (1.3%) |
| 1 | 25 (32%) |
| 2 | 31 (40%) |
| 3 | 13 (17%) |
| 4 | 4 (5.2%) |
| 5 | 3 (3.9%) |
| *Missing* | 1 |
| **pT** |  |
| *pT2a or 2b* | 4 (5.3%) |
| *pT2c* | 48 (64%) |
| *pT3a* | 16 (21%) |
| *pT3c* | 7 (9.3%) |
| *Missing* | 3 |

<sup>1</sup>Data are presented as the median value with inter-quartile range (IQR) in brackets or as number of values and percentage (%) in brackets PSA, prostate specific antigen

**Table S6:** List of primers used for RT-qPCR.

| **Gene name** | **Reference** |
| --- | --- |
| 18S | Hs99999901_s1 |
| CD36 | Hs00354519_m1 |
| FABP4 | Hs01086177_m1 |
| LIPE | Hs00943410_m1 |
| MGLL | Hs00996004_m1 |
| PNPLA2 | Hs00386101_m1 |
| SLC27A4 | Hs00192700_m1 |
| COL1A1 | Hs00164004_m1 |
| COL3A1 | Hs00943809_m1 |
| COL6A1 | Hs01095585_m1 |
| COL6A2 | Hs00365167_m1 |
| COL6A3 | Hs00915125_m1 |
| MMP14 | Hs00237119_m1 |
| MMP11 | Hs00968295_m1 |
| MMP2 | Hs01548727_m1 |
| MMP9 | Hs00957562_m1 |
| BMP1 | HS00241807_m1 |
| LOX | Hs00942483_m1 |
| LOXL1 | Hs00935937_m1 |
| TIMP1 | Hs99999139_m1 |
| TIMP2 | Hs00234278_m1 |
| TIMP3 | Hs00165949_m1 |
| TIMP4 | Hs00162784_m1 |

## References

1. Zwick RK, Guerrero-Juarez CF, Horsley V, Plikus MV. Anatomical, Physiological, and Functional Diversity of Adipose Tissue. Cell Metab. 2018;27:68–83.

2. Ishidoya S, Endoh M, Nakagawa H, Saito S, Arai Y. Novel anatomical findings of the prostatic gland and the surrounding capsular structures in the normal prostate. Tohoku J Exp Med. 2007;212:55–62.

3. Rosen ED, Spiegelman BM. What we talk about when we talk about fat. Cell. 2014;156:20–44.

4. Kahn CR, Wang G, Lee KY. Altered adipose tissue and adipocyte function in the pathogenesis of metabolic syndrome. J Clin Invest. 2019;129:3990–4000.

5. Estève D, Roumiguié M, Manceau C, Milhas D, Muller C. Periprostatic adipose tissue: A heavy player in prostate cancer progression. Current Opinion in Endocrine and Metabolic Research. 2020;10:29–35.

6. Nassar ZD, Aref AT, Miladinovic D, Mah CY, Raj GV, Hoy AJ, et al. Peri-prostatic adipose tissue: the metabolic microenvironment of prostate cancer. BJU Int. 2018;121 Suppl 3:9–21.

7. Passos GR, Ghezzi AC, Antunes E, de Oliveira MG, Mónica FZ. The Role of Periprostatic Adipose Tissue on Prostate Function in Vascular-Related Disorders. Front Pharmacol. 2021;12:626155.

8. Roumiguié M, Estève D, Manceau C, Toulet A, Gilleron J, Belles C, et al. Periprostatic Adipose Tissue Displays a Chronic Hypoxic State that Limits Its Expandability. Am J Pathol. 2022;192:926–42.

9. Miladinovic D, Cusick T, Mahon KL, Haynes A-M, Cortie CH, Meyer BJ, et al. Assessment of Periprostatic and Subcutaneous Adipose Tissue Lipolysis and Adipocyte Size from Men with Localized Prostate Cancer. Cancers (Basel). 2020;12:1385.

10. Woo S, Cho JY, Kim SY, Kim SH. Periprostatic fat thickness on MRI: correlation with Gleason score in prostate cancer. AJR Am J Roentgenol. 2015;204:W43–47.

11. Zhang Q, Sun L, Qi J, Yang Z, Huang T, Huo R. Periprostatic adiposity measured on magnetic resonance imaging correlates with prostate cancer aggressiveness. Urol J. 2014;11:1793–9.

12. Zhai T-S, Hu L-T, Ma W-G, Chen X, Luo M, Jin L, et al. Peri-prostatic adipose tissue measurements using MRI predict prostate cancer aggressiveness in men undergoing radical prostatectomy. J Endocrinol Invest. 2021;44:287–96.

13. van Roermund JGH, Hinnen KA, Tolman CJ, Bol GH, Witjes JA, Bosch JLHR, et al. Periprostatic fat correlates with tumour aggressiveness in prostate cancer patients. BJU Int. 2011;107:1775–9.

14. Liu B-H, Mao Y-H, Li X-Y, Luo R-X, Zhu W-A, Su H-B, et al. Measurements of peri-prostatic adipose tissue by MRI predict bone metastasis in patients with newly diagnosed prostate cancer. Front Oncol. 2024;14:1393650.

15. Xiong T, Cao F, Zhu G, Ye X, Cui Y, Xing N, et al. MRI-measured periprostatic adipose tissue volume as a prognostic predictor in prostate cancer patients undergoing laparoscopic radical prostatectomy. Adipocyte. 2023;12:2201964.

16. Jiang S, Li Y, Guo Y, Gong B, Wei C, Liu W, et al. MRI-measured periprostatic to subcutaneous adipose tissue thickness ratio as an independent risk factor in prostate cancer patients undergoing radical prostatectomy. Sci Rep. 2024;14:20896.

17. Dahran N, Szewczyk-Bieda M, Wei C, Vinnicombe S, Nabi G. Normalized periprostatic fat MRI measurements can predict prostate cancer aggressiveness in men undergoing radical prostatectomy for clinically localised disease. Sci Rep. 2017;7:4630.

18. Tan WP, Lin C, Chen M, Deane LA. Periprostatic Fat: A Risk Factor for Prostate Cancer? Urology. 2016;98:107–12.

19. Henriksen K, Genovese F, Reese-Petersen A, Audoly LP, Sun K, Karsdal MA, et al. Endotrophin, a Key Marker and Driver for Fibroinflammatory Disease. Endocr Rev. 2024;45:361–78.

20. Costello AJ. Considering the role of radical prostatectomy in 21st century prostate cancer care. Nat Rev Urol. 2020;17:177–88.

21. Pellegrinelli V, Carobbio S, Vidal-Puig A. Adipose tissue plasticity: how fat depots respond differently to pathophysiological cues. Diabetologia. 2016;59:1075–88.

22. Estève D, Boulet N, Belles C, Zakaroff-Girard A, Decaunes P, Briot A, et al. Lobular architecture of human adipose tissue defines the niche and fate of progenitor cells. Nat Commun. 2019;10:2549.

23. Ouchi N, Parker JL, Lugus JJ, Walsh K. Adipokines in inflammation and metabolic disease. Nat Rev Immunol. 2011;11:85–97.

24. Dupont S, Morsut L, Aragona M, Enzo E, Giulitti S, Cordenonsi M, et al. Role of YAP/TAZ in mechanotransduction. Nature. 2011;474:179–83.

25. Ma B, Cheng H, Gao R, Mu C, Chen L, Wu S, et al. Zyxin-Siah2-Lats2 axis mediates cooperation between Hippo and TGF-β signalling pathways. Nat Commun. 2016;7:11123.

26. Rausch V, Hansen CG. The Hippo Pathway, YAP/TAZ, and the Plasma Membrane. Trends Cell Biol. 2020;30:32–48.

27. Ichikawa T, Kita M, Matsui TS, Nagasato AI, Araki T, Chiang S-H, et al. Vinexin family (SORBS) proteins play different roles in stiffness-sensing and contractile force generation. J Cell Sci. 2017;130:3517–31.

28. Sun K, Tordjman J, Clément K, Scherer PE. Fibrosis and adipose tissue dysfunction. Cell Metab. 2013;18:470–7.

29. Hwang J, Huang Y, Burwell TJ, Peterson NC, Connor J, Weiss SJ, et al. In Situ Imaging of Tissue Remodeling with Collagen Hybridizing Peptides. ACS Nano. 2017;11:9825–35.

30. Bu D, Crewe C, Kusminski CM, Gordillo R, Ghaben AL, Kim M, et al. Human endotrophin as a driver of malignant tumor growth. JCI Insight. 2019;5:e125094, 125094.

31. Fujita K, Nonomura N. Urinary biomarkers of prostate cancer. Int J Urol. 2018;25:770–9.

32. Loeb S, Kettermann A, Carter HB, Ferrucci L, Metter EJ, Walsh PC. Prostate volume changes over time: results from the Baltimore Longitudinal Study of Aging. J Urol. 2009;182:1458–62.

33. Cescon M, Gattazzo F, Chen P, Bonaldo P. Collagen VI at a glance. J Cell Sci. 2015;128:3525–31.

34. Veidal SS, Karsdal MA, Vassiliadis E, Nawrocki A, Larsen MR, Nguyen QHT, et al. MMP mediated degradation of type VI collagen is highly associated with liver fibrosis--identification and validation of a novel biochemical marker assay. PLoS One. 2011;6:e24753.

35. Motrescu ER, Blaise S, Etique N, Messaddeq N, Chenard M-P, Stoll I, et al. Matrix metalloproteinase-11/stromelysin-3 exhibits collagenolytic function against collagen VI under normal and malignant conditions. Oncogene. 2008;27:6347–55.

36. Li X, Zhao Y, Chen C, Yang L, Lee H-H, Wang Z, et al. Critical Role of Matrix Metalloproteinase 14 in Adipose Tissue Remodeling during Obesity. Mol Cell Biol. 2020;40:e00564–19.

37. Jo W, Kim M, Oh J, Kim C-S, Park C, Yoon S, et al. MicroRNA-29 Ameliorates Fibro-Inflammation and Insulin Resistance in HIF1α-Deficient Obese Adipose Tissue by Inhibiting Endotrophin Generation. Diabetes. 2022;71:1746–62.

38. Tang Q, Cheng B, Dai R, Wang R. The Role of Androgen Receptor in Cross Talk Between Stromal Cells and Prostate Cancer Epithelial Cells. Front Cell Dev Biol. 2021;9:729498.

39. Karastergiou K, Smith SR, Greenberg AS, Fried SK. Sex differences in human adipose tissues – the biology of pear shape. Biol Sex Differ. 2012;3:13.

40. Sun K, Tordjman J, Clément K, Scherer PE. Fibrosis and adipose tissue dysfunction. Cell Metabolism. Cell Press; 2013;18:470–7.

41. Pellegrinelli V, Carobbio S, Vidal-Puig A. Adipose tissue plasticity: how fat depots respond differently to pathophysiological cues. Diabetologia. Springer Verlag; 2016;59:1075–88.

42. Khan T, Muise ES, Iyengar P, Wang ZV, Chandalia M, Abate N, et al. Metabolic dysregulation and adipose tissue fibrosis: role of collagen VI. Mol Cell Biol. 2009;29:1575–91.

43. Li X, Zhao Y, Chen C, Yang L, Lee H, Wang Z, et al. Critical Role of Matrix Metalloproteinase 14 in Adipose Tissue Remodeling during Obesity. Molecular and Cellular Biology. American Society for Microbiology; 2020;40.

44. Bonnans C, Chou J, Werb Z. Remodelling the extracellular matrix in development and disease. Nat Rev Mol Cell Biol. 2014;15:786–801.

45. Park J, Scherer PE. Adipocyte-derived endotrophin promotes malignant tumor progression. J Clin Invest. 2012;122:4243–56.

46. Olivares O, Mayers JR, Gouirand V, Torrence ME, Gicquel T, Borge L, et al. Collagen-derived proline promotes pancreatic ductal adenocarcinoma cell survival under nutrient limited conditions. Nat Commun. 2017;8:16031.

47. van Helvert S, Storm C, Friedl P. Mechanoreciprocity in cell migration. Nat Cell Biol. 2018;20:8– 20.

48. Dufau J, Shen JX, Couchet M, De Castro Barbosa T, Mejhert N, Massier L, et al. In vitro and ex vivo models of adipocytes. Am J Physiol Cell Physiol. 2021;320:C822–41.

49. Fontenot JJ, Lau FH. In Vitro Culture of White Adipose Tissue. Methods Mol Biol. 2024;2783:279–85.

50. Bouyssié D, Hesse A-M, Mouton-Barbosa E, Rompais M, Macron C, Carapito C, et al. Proline: an efficient and user-friendly software suite for large-scale proteomics. Bioinformatics. 2020;36:3148–55.

