## Supplemental figures for "Extra-cellular matrix remodeling as a unique mechanism of expansion of periprostatic adipose tissue: a potential driver of prostate cancer aggressiveness"

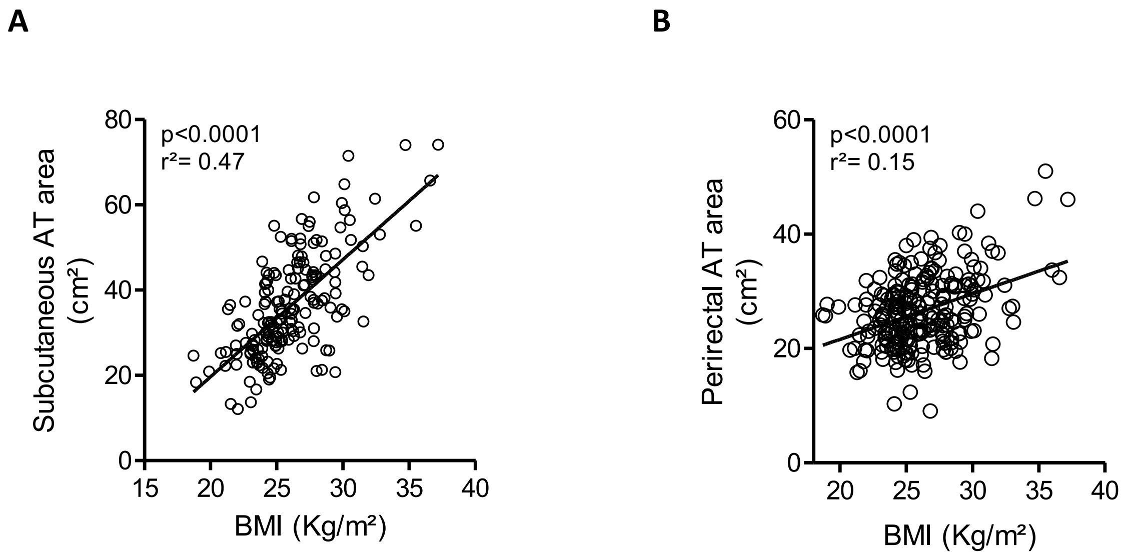


**Fig. S1: Subcutaneous and perirectal AT areas are correlated with BMI.** Linear regression between the subcutaneous (A, n= 195) and perirectal (B, n=256) adipose tissue area measured on pre-operative mpMRI and the BMI of the patients. Linear regression was used to draw the slope of best fit and linear correlation coefficient ($r^2$) and P-value are indicated (p).

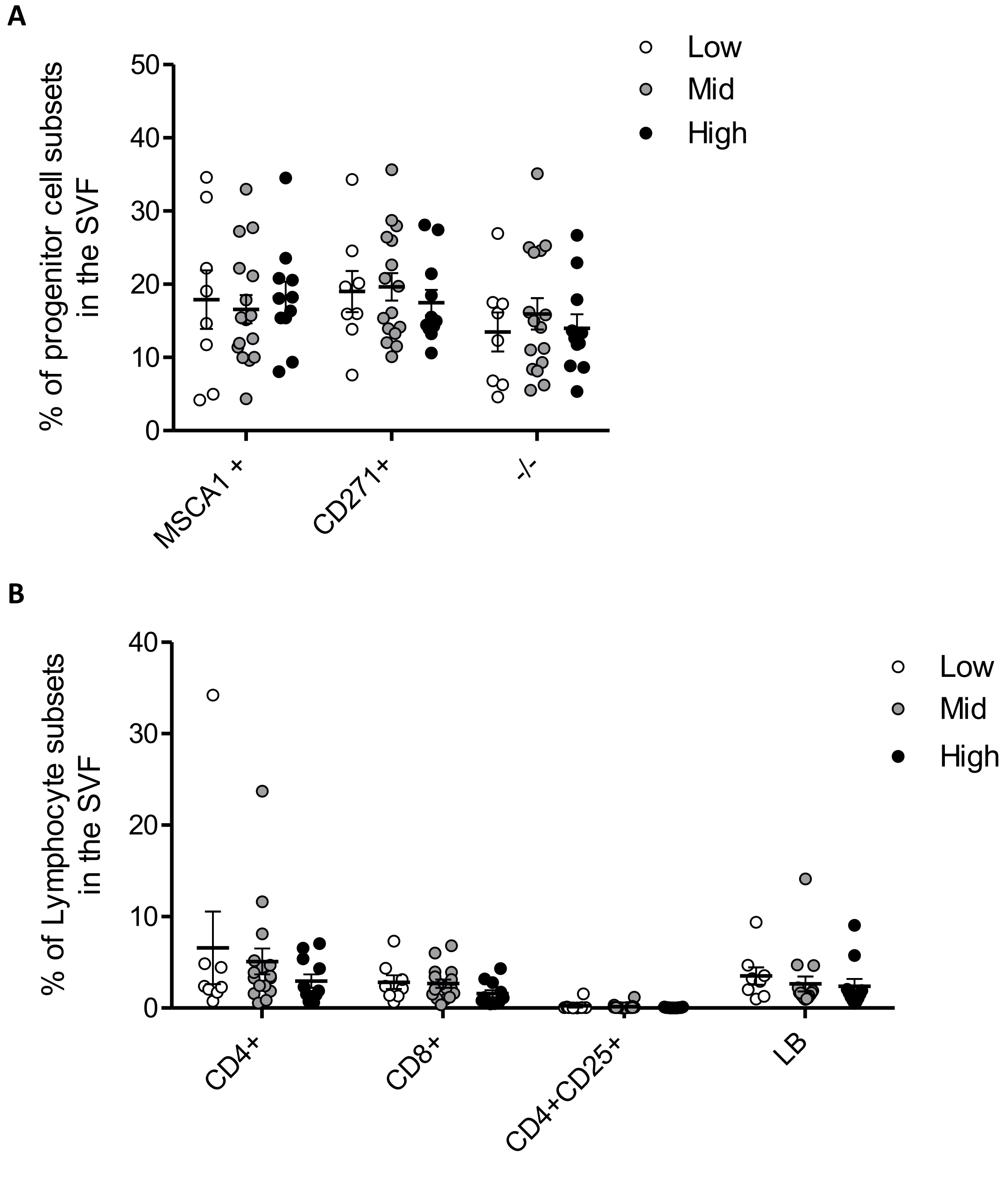


**Fig. S2: Adipose progenitor cell and lymphocytes subsets are not modified according to PPAT abundance. A)** Percentage of progenitor cell subsets (MSCA1+, MSCA1-/ CD271+ (CD271+) and MSCA1-/CD271- (-/-) in the stromal stromavascular fraction (SVF) of PPAT from patients in the low, mid and high groups, as quantified by flow cytometry. **B)** Percentage of lymphocytes subsets (CD4+: Cytotoxic T cell, CD8+: Helper T cell, CD4+/CD25+ cell, CD19+: B lymphocytes) gated on the total lymphocytes (Low FSC/Low SSC/CD45+/CD3+) in the stromal vascular fraction (SVF) of PPAT from patients in the low, mid and high groups, as quantified by flow cytometry. Bars indicate means ± SEM. No statistical differences were observed by using Student's T test.

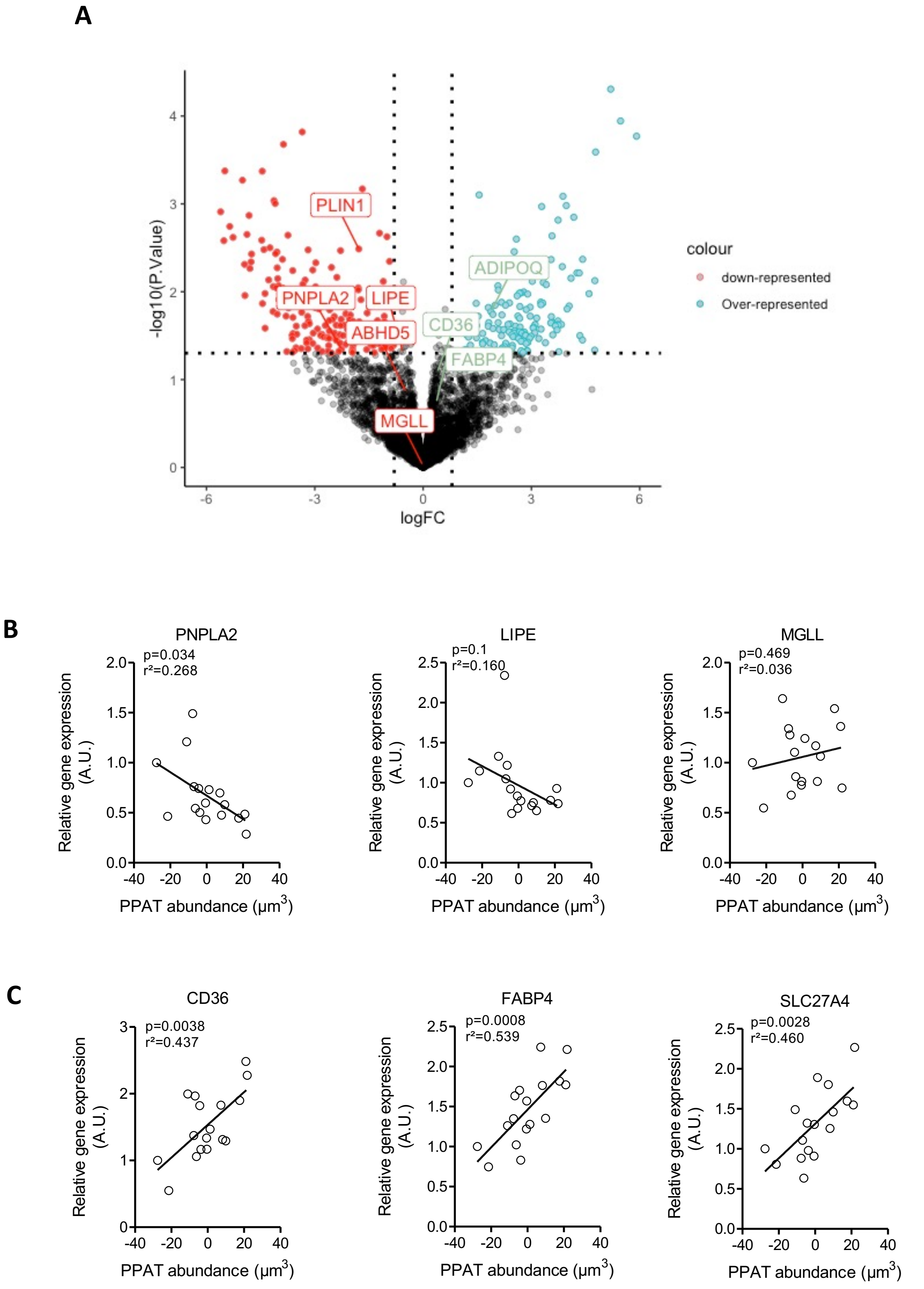


**Fig. S3: Proteins involved in lipolysis and lipid uptake are under-represented and over-represented in abundant PPAT. A)** Volcano plot representing the log2 fold-change of the relative protein representation in the proteomics dataset and the -log10 of the p-Value calculated by using limma differential expression analysis (n=6-7 /group), showing under-represented (red) and over-represented (green) proteins. Proteins of interest are indicated by name of their genes. **B**) Scatter plot of the relative expression of *PNPLA2*, *LIPE* and *MGLL*, which encode lipolysis effectors determined by RT- qPCR, according to PPAT abundance determined by our model defined in Figure 1D (n=18). **C)** Scatter plots of the relative gene expression of *CD36*, *FABP4* and *SLC27A4*, which encode lipid transporters determined by RT-qPCR, according to PPAT abundance determined by our model defined in Fig. 1D (n=18).

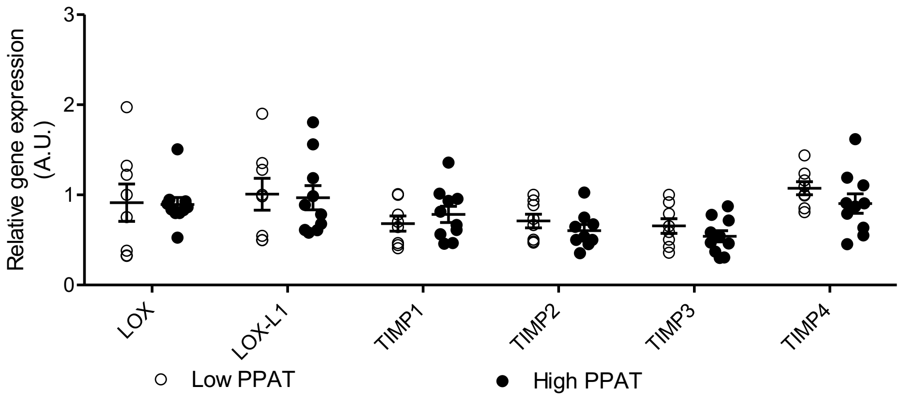


**Fig. S4: Absence of modifications of LOX or TIMP isoforms expression in low versus abundant PPAT.** Relative expression of several genes encoding lysyl oxidase (LOX) or tissue inhibitor of metalloproteinase (TIMP) isoforms in PPAT from patients in the low and high quartiles (n=11 per quartile). Bars indicate means ± SEM. No statistical differences were observed by using Student's T test.
